# Single amino acid substitution in DNA Polymerase I dramatically alters infection dynamics of bacteriophage T7

**DOI:** 10.64898/2026.05.20.726624

**Authors:** Rachel A. Keown, Andrew P. Sikkema, Victoria A. Barbone, Barbra D. Ferrell, Owen B. Donnelly, Sydney C. Iredell, Kelly M. Zatopek, Phillip J. Brumm, David A. Mead, Gregory J. S. Lohman, K. Eric Wommack, Shawn W. Polson

**Author notes:** deceased (Phillip J. Brumm).

## Abstract

Viruses constitute a significant proportion of Earth’s genetic diversity, yet most remain uncharacterized beyond their sequences in viral metagenomes. Linking viral genotypes to phenotypes—especially enzyme function to phage infection dynamics—is challenging due to the lack of cultured virus–host systems. DNA polymerase I (PolA), essential for genome replication in ∼25% of dsDNA phages, provides an opportunity to explore these connections. In phage T7, residue 526 is critical for nucleotide incorporation, with previous *in vitro* evidence indicating impacts on enzyme efficiency and fidelity. Previous analyses identified three substitutions at this position (Tyr/Y, Phe/F, Leu/L) linked with deeply rooted viral PolA clades. Mutation impacts at residue 526 were tested *in vitro* and *in vivo*. The Y526F protein exhibited a 50% reduction in specific activity, and when introduced via High Complexity Golden Gate Assembly into T7 demonstrated a 53% decrease in burst size and significantly longer latent period compared to wild type. The Y526L protein exhibited a 97% decrease in activity, and the Y526L phage was incapable of completing its lifecycle. These findings confirm historical biochemical data, provide *in vivo* context for these mutations in the T7–*E. coli* system, and offer experimental support for genotype-to-phenotype associations in viral PolA, informing viral metagenomics studies.

**GRAPHICAL ABSTRACT:** 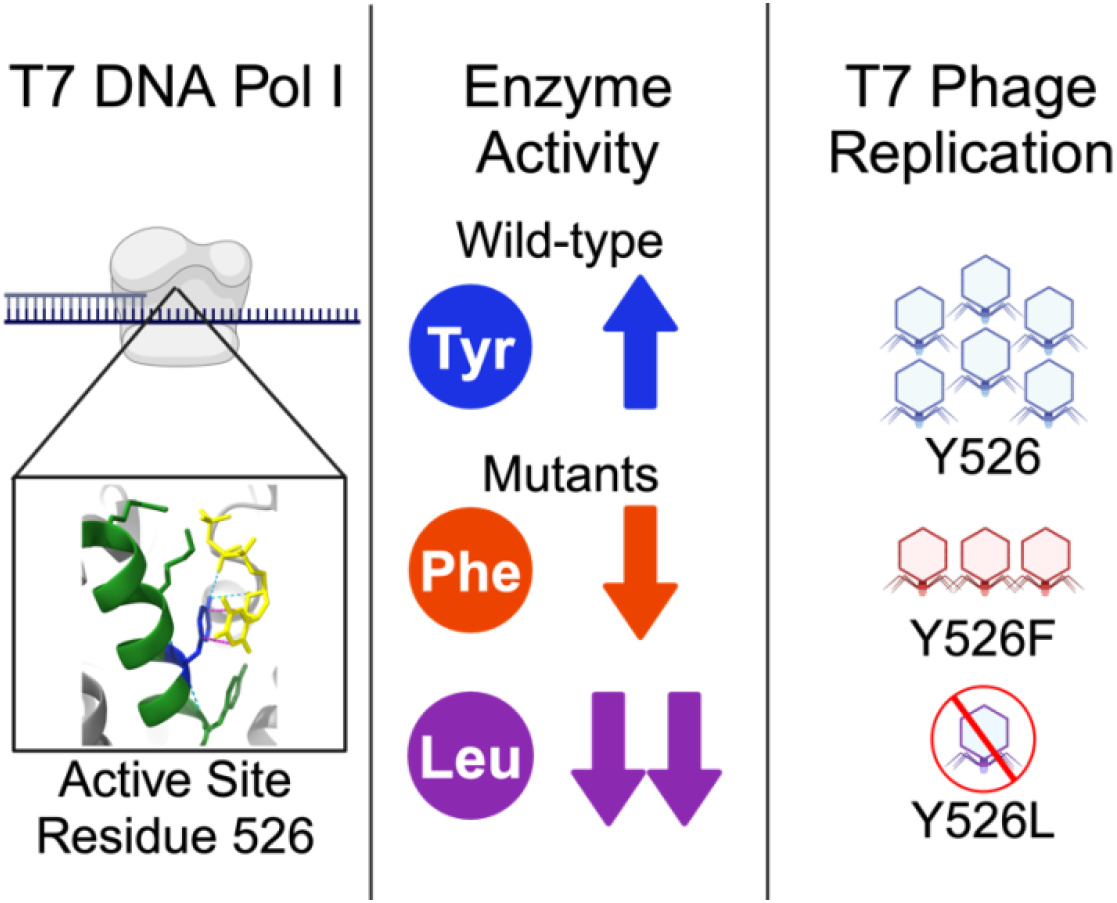

Created in BioRender. Keown, R. (2026) https://BioRender.com/mhrmup3

## INTRODUCTION

Metagenomics has uncovered the vast diversity of viruses within natural and host-associated microbial ecosystems (Rosario and Breitbart, 2011; Roux et al., 2021). Early work also found that viruses are extraordinarily abundant, oftentimes outnumbering co-occurring host microbes by 10 to 100-fold (Suttle, 2005) and accounting for a substantial amount of microbial mortality (Suttle, 2007). Thus, viral infection and lysis processes play important roles within ecosystems. Nevertheless, we still lack a fundamental understanding of the molecular mechanisms that govern viral infection dynamics and how these processes impact ecosystems. Enhanced understanding of genotype-to-phenotype connections within viruses could enable precise estimates of viral processes and their impacts on ecosystem function.

*Escherichia coli* phage T7 has long served as a model for interrogating the process of DNA replication (Richardson, 2015). The T7 replicon may be among nature’s simplest as only four proteins are required for phage genome replication: a single-stranded binding protein (gp2.5), DNA primase/helicase (gp4), DNA polymerase I (gp5), and host-associated thioredoxin (*E. coli trxA*) (Lee and Richardson, 2011). Metagenomic investigations have demonstrated that T7-like PolAs are common within natural viral populations (Breitbart and Rohwer, 2005; Schmidt et al., 2014, Nasko et al., 2018); thus, studies of the T7 model may be directly applicable for understanding ecosystem processes within viral communities. PolA is also common within bacteria serving as the DNA polymerase responsible for Okazaki fragment maturation (Makiela-Dzbenska et al., 2009). However, for viruses carrying PolA, the enzyme serves as the primary polymerase responsible for genome replication. Consequently, viral PolA enzymes exhibit expansive diversity demonstrating the enhanced selection pressure on this protein among viruses (Schmidt et al., 2014; Nasko et al., 2018; Keown et al., 2022).

The T7 DNA polymerase was developed as the first practical DNA sequencing enzyme (Zhu et al., 2014), and a great deal is known about how its *in vitro* biochemical characteristics connect with specific structural features. Position 526 was previously identified as critical for discriminating between chain-terminating ddNTPs and dNTPs (Tabor and Richardson, 1995). Specifically, the substitution of the wild-type tyrosine with phenylalanine reduced the discrimination between ddNTPs and dNTPs >1000-fold, as well as decreased the overall efficiency of the polymerase (Tabor and Richardson, 1995, Table 2). A leucine substitution at this site from a native phenylalanine in the *Thermus aquaticus* (Taq) PolA decreased nucleotide incorporation efficiency ten-fold but increased the overall base selection fidelity of the polymerase (Suzuki et al., 2000).

Studies of DNA polymerase I sequences have shown that all three residue variants (tyrosine, phenylalanine, and leucine) are observed in unknown viruses at the position homologous to T7 DNA PolA 526 (762 in *E. coli* DNA PolA) (Schmidt et al., 2014; Nasko et al., 2018; Keown et al., 2022). These studies have hypothesized that because this position has such dramatic effects on PolA biochemistry it may connect with viral life history strategy. Schmidt et al. (2014) noted that tyrosine occurred almost exclusively in known virulent phage (i.e. phage exclusively undergoing a lytic lifecycle), phenylalanine was common in virulent phage, and leucine occurred predominantly in temperate phage (i.e. phage capable of a lysogenic lifecycle).

While there is a wealth of *in vitro* biochemical and evolutionary data, the *in vivo* impacts of 526 mutations on phage infection dynamics are unknown, even though these mutations commonly occur in nature. Addressing this knowledge gap using the T7–*E. coli* phage–host system was the central objective of this investigation. One step assembly of T7 genomes was recently accomplished through High Complexity Golden Gate Assembly (HC-GGA) (Parker et al., 2025; Sikkema et al., 2023; Pryor et al., 2020 and 2022). Here GGA was used for introducing point mutations within the T7 DNA polymerase in the context of the full phage genome. Both *in vitro* and *in vivo* approaches were used for assessing enzyme activity and impacts on phage phenotype. By connecting PolA genotypic changes to changes in infection phenotype, this study marks a significant step forward in uncovering possible molecular mechanisms underlying phage infection dynamics within nature.

## MATERIAL AND METHODS

### Protein purification

Mutant DNA polymerases (PolAs) were generated via site-directed mutagenesis of wild-type T7 genomes (mutational primers in Supplementary Table S1, denoted with **). Mutant 526 PolAs were assembled into a co-expression vector with wild-type *E. coli* thioredoxin (*trxA*), generated via PCR (primers in Supplementary Table S1). Both genes were cloned between *Bam*HI and *Not*I sites of an arabinose-inducible pD871-based vector containing an N-terminal 9xHis tag (Chiu et al., 2006; plasmid map in Supplementary Fig. S1). TrxA co-expression and 9xHis tagging produced higher levels of soluble protein as described by Chiu et al. (2006). Plasmid assemblies were chemically transformed into *E. coli* 5-α cells (New England Biolabs, Ipswitch, MA) and plated on selective media (Luria-Bertani (Lennox, 1955), supplemented with 30 µg/mL kanamycin, 0.4% (w/v) dextrose). Single colonies were isolated and grown for plasmid purification and 50% (v/v) glycerol stock storage. Whole plasmids were sequence confirmed with Oxford Nanopore sequencing (SNPsaurus LLC, Eugene, OR).

Cultures for protein purification were grown as described in Keown et al., 2022, with minor modifications. Briefly, protein products from the collected fractions of the HisPur™ Ni-NTA Resin (ThermoFisher Scientific, Waltham, MA) column containing the protein of interest were pooled, diluted 1:1 in molecular grade water, and loaded onto a 20 mL Q Sepharose (Cytiva, Marlborough, MA) column pre-equilibrated with extraction buffer (50 mM Tris-HCl pH 8, 125 mM NaCl, 15 mM imidazole). The column was washed with ten volumes of extraction buffer, and the polymerase was eluted with ten column volumes of elution buffer (100 mM Tris-HCl pH 8, 250 mM NaCl, 30 mM imidazole) and collected in 6 mL fractions.

The eluent was concentrated by dialysis with storage buffer A (50 mM Tris-HCl pH 7.5, 50 mM KCl, 1 mM DTT, 1 mM EDTA, 50% glycerol) creating DNA polymerase protein stocks. The dialyzed protein product was confirmed via SDS-PAGE using 4–20% Mini-PROTEAN TGX precast protein gels (Bio-Rad Laboratories, Hercules, CA). Gels were stained with Pierce™ 6xHis Protein Tag Stain Reagents (ThermoFisher Scientific) following the manufacturer’s protocol and 0.1% Coomassie R-250 (Research Products International, Mount Prospect, IL), 40% ethanol, 10% acetic acid, followed by destaining, and imaged. Exemplary dual-stained PAGE gels are shown in Supplementary Fig. S2. T7 DNA polymerase protein stocks were quantitated by measuring absorbance at 280 nm and subsequently used in all downstream enzyme assays.

### Primer extension assays

Primer extension assays were performed in 1X T7 DNA polymerase reaction buffer (40 mM Tris-HCl pH 7.5, 10 mM MgCl2, 1 mM DTT) as described previously (Keown et al., 2022). Reactions (30 µL) were isothermally incubated at various temperatures for 2 h, then mixed with 6X loading dye with SDS (ThermoFisher Scientific) and heat-killed for 2 min at 70 °C. Products were visualized on a 0.7% agarose gel in 1X TAE buffer with 1X SYBR Safe stain (ThermoFisher Scientific) run at 80 V for 1 h. Positive (double-stranded) products were qualitatively placed into categories of strong, moderate, or weak activity.

### Specific activity assays

Specific activity assays were performed in 1X T7 DNA polymerase reaction buffer. Twenty-five microliter reactions included 1 µg of M13mp18 single-stranded phage DNA (Bayou Biolabs, Metairie, LA), 3.3 µM ssM13 FT primer with three phosphorothioate bonds (*) introduced at the 3′ end (5′-CGCCAGGGTTTTCCCAGTCAC*G*A*C-3′ Integrated DNA Technologies, Coralville, IA), 200 µM dNTPs (New England Biolabs), 1X SYBR Green I (ThermoFisher Scientific) and seven 1:1 serial dilutions of an initial 5 µg WT T7 DNA polymerase (or variant)/reaction concentration in 1X sample buffer. Reactions were incubated at 37 °C for 10 min using a Bio-Rad Laboratories CFX96 Real-Time PCR machine. Fluorescence was quantified in the SYBR channel (497/520 nm) every 3 sec for 40 cycles. Data was analyzed by manual RFU background subtraction and slope determination of the initial cycles representing enzyme activity. Slopes were plotted against the standard curve (RFU/cycle vs enzyme units) of the average of three technical replicates of a control polymerase (commercially available T7 DNA polymerase, ThermoFisher Scientific, Cat# EP0081; one unit of the enzyme catalyzes the incorporation of 10 nmol of deoxyribonucleotides into a polynucleotide fraction in 30 min at 37 °C) to convert values to Units of enzyme activity. Units per microgram and Units per microliter were calculated based on the test enzyme’s quantity per reaction (µg) and stock concentration (µg/µL). Reported values are the average of three technical replicates. Statistical significance was determined using a pairwise t-test with Bonferroni correction in R (v4.3.0) using the ‘stats’ package (v4.3.0). A p-value less than 0.05 was considered statistically significant.

### Fidelity assays

To accurately measure the fidelity of T7 DNAP variants, we employed a Pacific Biosciences (PacBio) single-molecule assay (Pacific Biosciences, Menlo Park, CA). The 2kb template DNA utilized for fidelity measurements was synthesized as previously described (Betancurt-Anzola et al., 2025). Primer extension of the 2 kb ssDNA template by the three purified mutant T7 DNA polymerases was performed by annealing 5 µL of the forward primer (5′-GGGAAGCAGACGTAATATATG-3′) at 100 µM to 35 µL of the 2 kb ssDNA by incubation at 65 °C for 5 min followed by transfer to ice. Primer extension was carried out in a 50 µL reaction by incubating 5 µL of T7 10X reaction buffer (400 mM Tris-HCl, 100 mM MgCl_2_, 10 mM DTT) containing 200 µM dNTPs (New England Biolabs), 50 nM mutant DNA polymerase, and 40 µL of annealed substrate at 37 °C for 1 h. Extended products were cleaned using NEB sample purification beads and eluted in 50 µL low TE buffer. Second strand synthesis was carried out in triplicate for each mutant T7 DNA polymerase.

PacBio sequencing: PacBio libraries were created from the extension products utilizing the PacBio SMRTbell prep kit 3.0 (Pacific Biosciences, Menlo Park, CA) following the manufacturer’s protocol: “Preparing multiplexed amplicon libraries using SMRTbell prepkit 3.0” starting on step 3 of the protocol and barcoding utilizing the SMRTbell Barcoded Adapter Plate 3.0. Note that, on step 3.1, the DNA repair mix was replaced with nuclease free water.

The PacBio SMRTlink was used to prepare samples for sequencing utilizing the <3 kb Amplicons method, the Sequel® II Binding Kit 3.1, and 150 pM concentration on plate. The sequencing run for Sequel® II was set up with default parameters for the <3 kb Amplicons application and 20 h of movie time. Following sequencing, data were analyzed to extract sequencing errors as previously described (Betancurt-Anzola et al., 2025).

### T7 genome 13-fragment assembly design

The 13-part T7 Golden Gate Assembly was designed following bacteriophage genome assembly principles outlined previously (Sikkema et al., 2023). The previously reported BsmBI domesticated T7 genome (generated by Pryor et al., 2022) was used as a starting point (see Supplementary Table S1 for primer sequences). Using an already domesticated genome allowed the fusion sites between fragments to be placed without the need for mutagenizing pre-existing Type IIS recognition sites in the T7 genome. Twelve fusion sites were selected creating a modular genome with fragments of convenient size for PCR with a high-fidelity junction set among them that would permit the simple addition of more breakpoints without compromising fidelity. An additional 13^th^ fusion site was placed in fragment 7 near Y526 of the DNA polymerase gene (gp5), resulting in fragments 7a and 7b, which were used for introducing mutations via PCR primers following the same procedure used to place domesticating mutations in Protocol 5 of Sikkema et al., 2023. See Supplementary Fig. S3 for genome map and Supplementary Table S2 for breakpoints.

### ΔDNAP assembly design

To remove the DNA polymerase gene (gp5) from the T7 genome, fragments 6 and 7, from the starting 12-part assembly, were combined then broken down into smaller fragments, one of which contained the DNA polymerase gene (gp5). Adjacent fragments were generated by PCR (see below), while the new DNA pol fragment was designed *in silico*, replacing the coding region of gp5 with sfGFP, leaving the start and stop codons intact (Supplementary Fig. S3 and Supplementary Table S2). M13/pUC primer sites were added to the ends of the fragment, and the final fragment was ordered from Twist Biosciences (San Francisco, CA) as a Twist Fragment (Supplementary Table S3). The Twist Fragment was used directly for the assembly reaction with the M13/pUC primer sites allowing for additional input DNA to be made by PCR if needed.

### T7 genome fragment generation

All phage T7 genome fragments were generated via PCR using Q5 High Fidelity 2X PCR Master Mix (New England Biolabs). Primer sequences are listed in Supplementary Table S1. Triplicate 50 µL PCR reactions were performed as follows: 1 ng of BsmBl domesticated T7 phage gDNA (Pryor et al., 2022), 0.5 µM forward and reverse primers, 25 µL Q5 master mix, and 19 µL molecular grade water. PCR conditions for each fragment, including primer names, annealing temperatures, and fragment lengths, are listed in Supplementary Table S2. Cycling conditions were as follows: 30 sec at 98 °C as initial denaturation, followed by 35 cycles of 10 sec at 98 °C for denaturation, 3 min at variable annealing temperature (see Supplementary Table S2), 40 sec at 72 °C for extension, and final extension at 72 °C for 2 min.

PCR reactions were then pooled and concentrated using the Monarch PCR clean up kit with one additional spin for removing residual ethanol prior to elution. Cleaned products were quantified using Qubit dsDNA HS kit.

### Golden Gate assembly of phage genomes

HC-GGA fragments (PCR products) were combined into an equimolar master mix of each fragment. Thirteen-fragment assemblies containing position 526 mutations were designated as follows: Y526Y wild-type tyrosine assembled without mutagenesis; Y526F phenylalanine substitution assembled with site-specific mutagenesis; and Y526L leucine substitution assembled with site-specific mutagenesis; ΔDNAP refers to the polymerase deletion mutant, assembled as described above. High Complexity Golden Gate assemblies were performed in 0.5X reactions using the NEBridge Golden Gate Assembly Kit (BsmBI-v2) (New England Biolabs) per the manufacturer’s protocol at a final concentration of 3 nM of each fragment. Thermal cycling conditions were as follows: 90 cycles of 5 min at 42 °C for overhang generation, and 5 min at 16 °C for ligation, followed by a final incubation of 5 min at 60 °C; lid temp 75 °C. Assemblies were verified via agarose gel electrophoresis and sequence-confirmed with Illumina sequencing (SeqCenter, Pittsburgh, PA), long-read sequencing (Oxford Nanopore Technologies, Oxford, United Kingdom), and capillary electrophoresis (Agilent TapeStation, Santa Clara, CA) analysis (Sikkema et al., 2023, Support Protocol 4).

### Generation of PolA rescue strain electrocompetent cells

An untagged, codon-shuffled T7 DNA polymerase gene (gp5) was inserted into a custom pET backbone that lacked the LacO site. Codon shuffling ensured that genome repair through crossover or recombination was not possible. This design placed the *polA* gene under the control of the T7 promoter (Supplementary Fig. S4, Supplementary File S1). Plasmids were transformed into NEB 10β cells (New England Biolabs). One half liter of LB media with 50 µg/mL kanamycin was inoculated with 5 mL of stationary phase LB 50 µg/mL kanamycin overnight culture of NEB 10β cells carrying the wild-type T7 PolA plasmid, and grown at 37 °C, 225 rpm until OD600 reached ∼0.6. Cells were harvested by centrifugation at 4000 x g for 15 min. The supernatant was decanted, and residual media was removed using a serological pipette. The cell pellet was washed three times by completely resuspending in 300 mL ice-cold 10% glycerol and pelleting at 4000 x g for 10 min. After pelleting the third wash, the supernatant was removed using a serological pipette and the cells were resuspended in 500 µL ice-cold 10% glycerol to generate a 1000X concentrated stock. PolA rescue electrocompetent cell aliquots (50 µL) were frozen in a dry ice ethanol bath and stored at −80 °C.

### Transformation of assembled genomes

Assemblies were transformed into NEB 10β electrocompetent cells (New England Biolabs) or PolA rescue electrocompetent cells using Bio-Rad Laboratories GenePulser with 1800 kV preset program (Ec1; Bio-Rad Laboratories) in 1 mm electroporation cuvettes (BTX Model 610: 45-0124; ThermoFisher Scientific). Cells were recovered on SOC media or NEB 10-beta/Stable Outgrowth Medium (New England Biolabs) at 37 °C and 225 rpm for 1 h. Cells were plated on Luria-Bertani (LB 1.5% agar underlay, 0.7% agar overlay) plates and incubated at 37 °C for 4 h or until plaques developed (overnight). Single plaque isolates were sequence-confirmed via plaque isolation, PCR amplification, and Sanger sequencing of a 500 bp fragment of the DNA polymerase gene (University of Delaware DNA Sequencing and Genotyping Center, Newark, DE; primers in Supplementary Table S1). Plates were imaged using Bio-Rad Laboratories Gel Doc XR+ with settings: standard filter, blue transillumination, exposure for faint bands, image color grey.

### Viral lysate preparation

*E. coli* B cultures were infected with sequence-verified single plaque 526 mutant T7 isolates and plated on LB agar media. Plaques were allowed to overdevelop and clear all bacterial growth at 37 °C overnight. Once cleared, plates were flooded with MSM buffer (50 mM Tris HCl, 400 mM NaCl, 20 mM MgSO_4_) and incubated with rocking for 1 h at room temperature. After incubation, buffer was aspirated from the plate surface and collected into 50 mL conical tubes. Top agar was then scraped from the plates and added to a 50 mL conical tube with 25 mL fresh MSM buffer and vortexed for 5 min at high speed to break up any large pieces. All tubes were centrifuged at 4000 x g for 5 min to pellet any solids. The supernatant was removed and centrifuged twice more. The cleaned supernatant was then filtered through 0.2 μM Whatman syringe top filters (GE Healthcare, Chicago, IL) removing bacterial cells and agar fragments. Filtered supernatants were concentrated using 100 kDa Amicon filters (Sigma-Aldrich, Saint Louis, MO). Cleaned and concentrated viral lysates were titered against *E. coli* B cells and concentrations were noted in plaque-forming units per milliliter (PFU/mL). Viral lysates were used in all subsequent phenotyping experiments.

### One-Step Growth Curve

Phage phenotyping was accomplished using the one-step growth curve experiment (Kropinski, 2018). *E. coli* B cells were cultured in LB broth at 37 °C. Cultures were diluted 1:100 and grown until the OD600 reached 0.3–0.4 and cells were in a logarithmic growth phase, approximately 4 h post culture initiation. Cultures were then infected with titered phage stocks (viral lysates) at a multiplicity of infection (MOI) of 0.01. Phages were allowed to adsorb to cells for 2 min at 37 °C, and then cells were pelleted by centrifugation (4000 x g for 5 min) to remove unadsorbed phages. Cell pellets were resuspended in fresh LB warmed to 37 °C. Once resuspended, sampling began at T_0_ and continued every ∼2 min for 1 h. At each sampling, optical density (OD600) was measured and 45 µL samples were diluted 1:100 in LB and plated in triplicate for PFU counts. The gDNA phage strain included was a methodological control to ascertain if HC-GGA had any impact on latent period or burst size. Latent period was calculated from the initial time of infection, including resuspension time. Burst size was calculated from the average PFU in the constant period minus the average PFU in the latent period, divided by the number of cells for a measure of PFU per infected cell.

### T7 genome replication qPCR assay

Quantitative PCR was performed on all T7 phage PolA mutant strains after transformation into NEB 10β cells. Cells were allowed to recover after electroporation as described above, and 500 µL of outgrowth was subsequently used to inoculate 19.5 mL of LB broth equilibrated to 37 °C. Cultures were grown at 37 °C for 32 h and sampled at regular timepoints. At each sampling time, OD600 was measured and 200 µL was retained, snap-frozen in liquid nitrogen and stored at −80 °C prior to qPCR analysis.

Two regions of the T7 genome, a 103 bp region in the RNA polymerase gene, and a 101 bp region in the terminase large subunit gene (primers in Supplementary Table S1) were targeted for quantification. Each region was at opposite ends of the genome enabling testing for full-length genome replication. All samples and standards were performed in triplicate using Luna® Universal qPCR Master Mix (New England Biolabs). Samples were prepared and cycling conditions were performed per the manufacturer’s protocol (New England Biolabs) for 40 cycles on the Bio-Rad Laboratories CFX96 Real-Time PCR machine.

### Alignment and Phylogeny

Environmental PolA sequences were retrieved and validated as described in Keown et al. (2022). These data included bacterial and viral reference sequences, and 675 sequences from the Global Ocean Virome (GOV) study. Sequences were aligned and used for approximate maximum likelihood tree building as described previously. Branches were colored to denote 762 (*E. coli*) position identity using Iroki (Moore et al., 2020).

## RESULTS

### Position 526 mutations in bacteriophage T7 DNA polymerase decrease efficiency but maintain fidelity

Prior work had shown that co-purification of WT and mutant PolA with *E. coli* thioredoxin was necessary for full and consistent enzyme activity (Mark and Richardson, 1976; Tabor and Richardson, 1995); a strategy which we also employed to purify our T7 PolA proteins (Supplementary Fig. S1 and S2). Primer extension activity varied across temperature and concentration gradients between the wild-type and mutant enzymes (Supplementary Fig. S5). For example, at the highest protein concentration (5 µg/rxn) the wild-type (Y526Y) enzyme displayed strong activity at all temperatures (5–54°C), the phenylalanine mutant (Y526F) enzyme showed a narrower range of strong activity (5–41°C), and the leucine mutant (Y526L) showed only weak primer activity at 37 °C. Follow-up quantitative *in vitro* primer extension assays showed a 50% decrease in specific activity for the Y526F mutant and a 97% decrease for the Y526L mutant compared to the wild-type enzyme (Y526Y) (Fig. 1). These decreases were statistically significant (P < 10^-10^).

**Figure 1.**
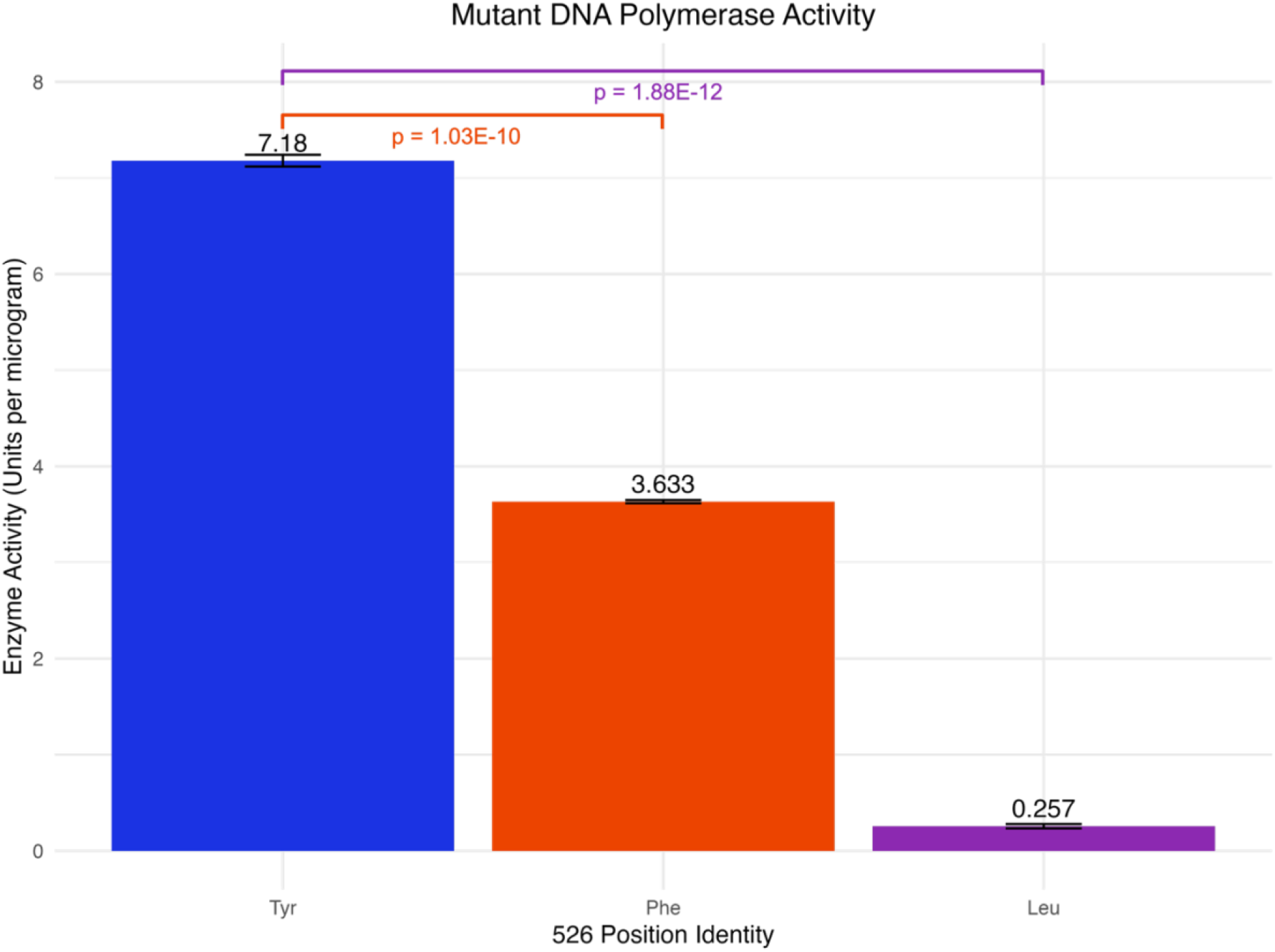
Mutation of amino acid 526 significantly decreases *in vitro* polymerase activity. Normalized specific activity for polymerase stocks presented in Units per microgram of enzyme as compared to commercially available T7 DNA polymerase (ThermoFisher Scientific; one unit of the enzyme catalyzes the incorporation of 10 nmol of deoxyribonucleotides into a polynucleotide fraction in 30 min at 37 °C). Triplicate reactions were performed in 1X T7 reaction buffer with ssDNA, forward primer, dNTPs, and 1X SYBR Green I to measure real-time dNTP incorporation via fluorescence. The Y526F mutant enzyme showed a 50% reduction in speed, while Y526L showed a 97% reduction as compared to wild-type tyrosine. Significance according to pairwise t-tests with Bonferroni correction is indicated by brackets between bars. Created in BioRender. Keown, R. (2026) https://BioRender.com/7j2e7gr

Enzyme fidelity assays were undertaken for all PolA variants but successfully measured for only the wild-type (Y526Y) and mutant Y526F. The Y526L protein did not successfully complete second strand synthesis of the 2 kb template and no sequencing library could be produced. No significant differences in PolA fidelity (p = 0.267) were found between the Y526Y and Y526F enzymes (Supplementary Fig. S6). Substitution spectra show the most common transversion was from thymine to adenine for both enzymes (Supplementary Fig. S7), in line with the expected error profile for DNA polymerase I enzymes (Suzuki et al., 2000).

### Generation of mutant T7 bacteriophage genomes and phage boot up

Golden Gate Assembly enabled precise single amino acid substitutions at position 526, generating three mutant T7 phage strains: Y526Y (wild-type control), Y526F, and Y526L. The genomes were assembled *in vitro* and validated using both capillary electrophoresis and whole-genome sequencing (Supplementary Fig. S8). After assembly, mutant T7 genomes were transformed into *E. coli* 10β cells. The Y526Y and Y526F genomes formed plaques under standard conditions (Fig. 2A and C). These plaques were sequence-verified, propagated, and purified producing high-titer virus lysate stocks. The Y526L genome failed to produce plaques under standard conditions (Fig. 2E) and a specialized *E. coli* rescue strain was required (see below). Rescued plaques were isolated, and the identity of the 526 position was confirmed with Sanger sequencing.

**Figure 2.**
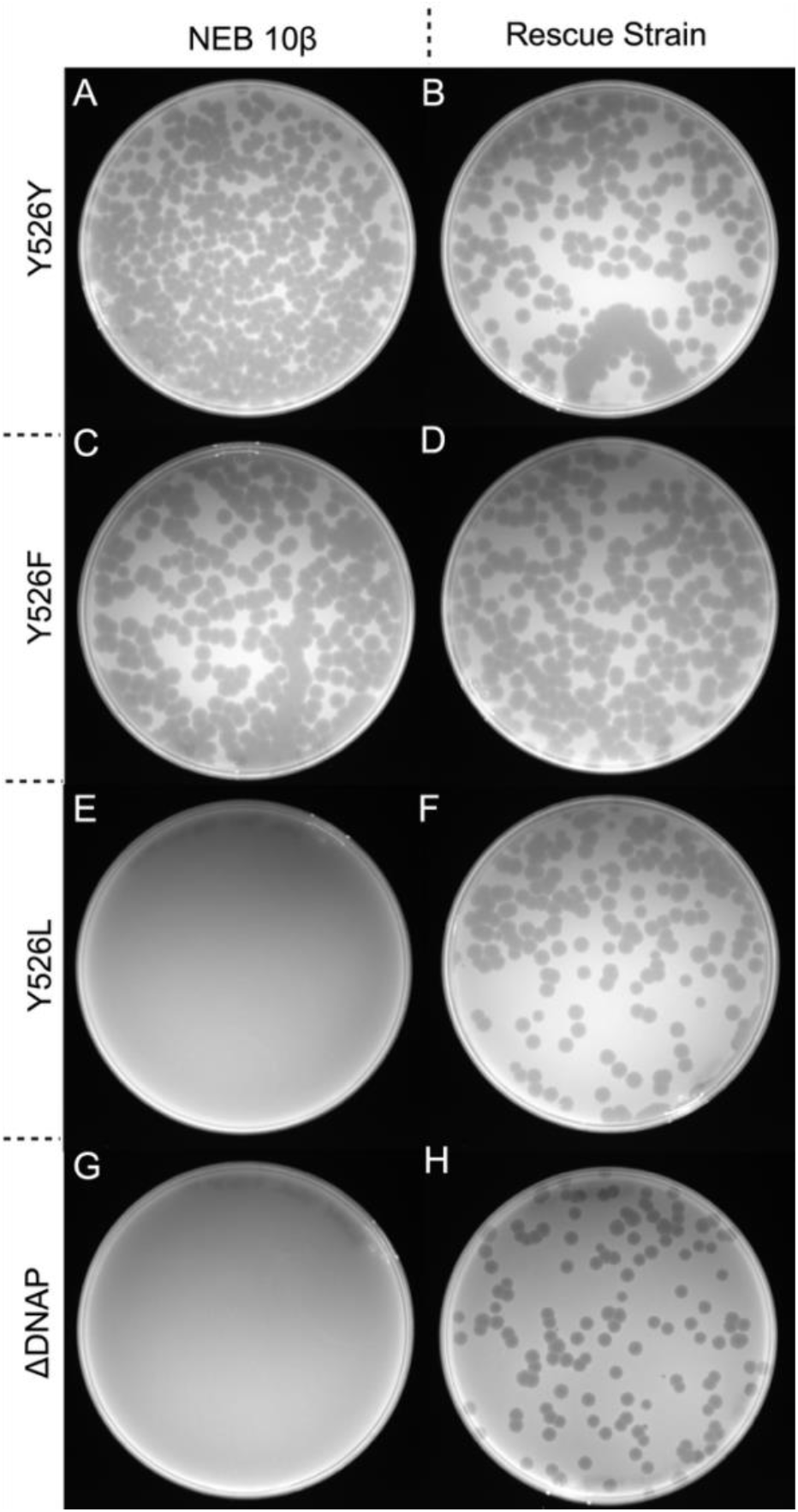
Plaque assays suggest Y526L point mutation interrupts phage life cycle. Plaque assay of transformed *E. coli* 10β wild-type (left) and rescue strain (right), containing a codon-shuffled wild-type T7 DNA *polA* gene on a plasmid. Cells were transformed, recovered, and plated; incubated at 37 °C for 8 h for plaque and lawn to develop; and imaged. Wild-type phage form plaques on *E. coli* A) wild-type and B) rescue strains. Mutant Y526F phage form plaques on *E. coli* C) wild-type and D) rescue strains. Mutant Y526L E) does not form plaques on wild-type *E. coli*, but F) does form plaques on rescue cells. The negative control ΔDNAP phage shows a rescued phenotype similar to Y526L in that it G) does not form plaques on wild-type *E. coli*, but H) does form plaques on rescue cells. These results indicate that the Y526L point mutation disrupts the phage life cycle in wild-type cells, but this phenotype can be rescued in the presence of wild-type PolA expressed in the rescue strain. Sequencing of recovered phage plaques confirm the presence of the Y526L within mutant phages grown on the rescue strain. Created in BioRender. Keown, R. (2026) https://BioRender.com/cyfggqe

### PolA position 526 mutations substantially alter T7 infection phenotypes

Infection phenotypes of three recombinant T7 phage genomes (Y526Y, Y526F, and Y526L) and a PolA knockout genome (ΔDNAP, replacing PolA with sfGFP) were assessed. The ΔDNAP T7 phage genome was designed to require replication in host cells containing a PolA gene (Fig. S3) to mimic genomes containing impaired PolA. A host strain carrying a plasmid-encoded PolA under the control of a T7 promoter (Supplementary Fig. S4) was generated to provide the necessary replication machinery to allow PolA-deficient strains to replicate. The plasmid-encoded *polA* gene was codon-shuffled to prevent genome repair by crossover or recombination. The Y526L and ΔDNAP genomes failed to form plaques upon transformation into NEB 10β cells but did produce plaques on the rescue strain (Fig. 2E–H). The Y526Y and Y526F mutants showed comparable plaque-forming activity on both NEB 10β and PolA rescue strains (Fig. 2A–D). Ong et al. (2026) used a similar strategy of a host strain containing a plasmid-encoded *polA* gene for replication of an engineered T7 lacking a *polA* gene as part of an *in vivo* evolution system.

Quantitative PCR (qPCR) was performed on T7 GGA transformations into NEB 10β cells for determining whether some level of phage replication occurred with Y526L. Assays targeted the RNA polymerase and terminase (large subunit) genes. Log-fold increases in both gene targets were observed for the Y526Y and Y526F phage transformants, confirming the success of these polymerases in T7 genome replication (Supplementary Fig. S9A). In contrast, minimal change in qPCR signal was observed for the Y526L T7 transformation over a 2 h incubation (Supplementary Fig. S9A). All assemblies remained quantifiable in liquid culture over 32 h (data not shown). OD600 measurements showed the opposite trend to phage DNA copy number, increasing in both the negative control (ΔDNAP) and the Y526L mutant and decreasing in the Y526Y and Y526F phage transformations (Supplementary Fig. S9B).

### The Y526F mutation impacts viral infection dynamics

One-step growth curve assays found that the Y526Y phage showed no statistically significant difference in latent period or burst size as compared to the gDNA (wild-type) control infection (Fig. 3). In contrast, the Y526F phage showed a 15% increase in latent period (p = 1.19E-06) and a 53% decrease in burst size (p = 2.67E-09) as compared to the Y526Y phage. The Y526L phage did not form plaques on wild-type *E. coli* (Fig. 2) and did not show evidence of *in vivo* replication in the qPCR assay (Supplementary Fig. S9). It is interesting to note that qPCR findings also supported these phenotypic changes in infection dynamics phenotypes between Y526Y and Y526F (Supplementary Fig. S9). Rises in T7 genome copy number occurred later in the Y526F mutant as compared with Y526Y. Similarly, reductions in host cell abundance (assessed from OD600) began decreasing later for infections with the Y526F phage mutant.

**Figure 3.**
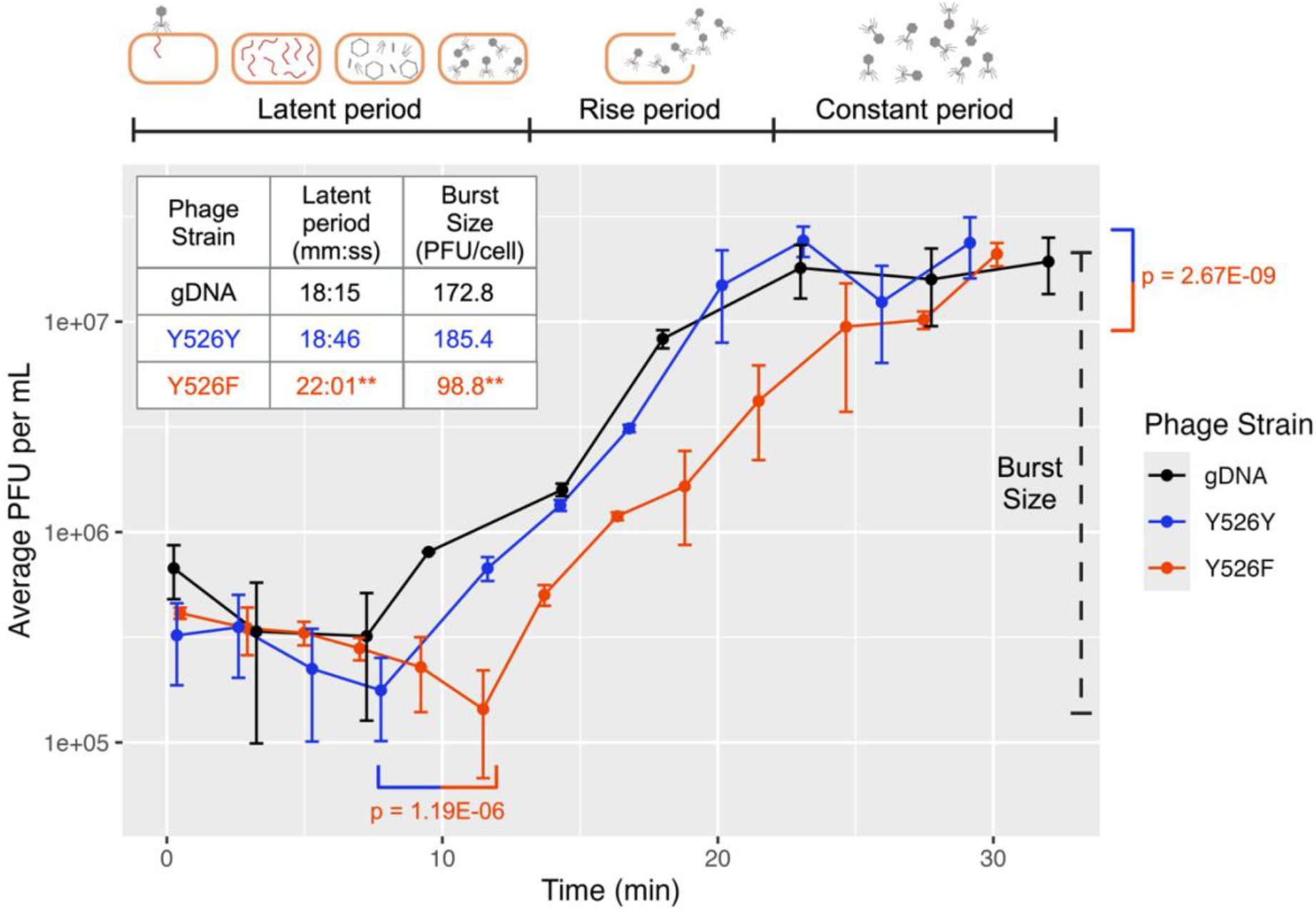
Altered Infection Dynamics of T7 phages. One-step growth curve (OSGC) of control (gDNA) and High Complexity Golden Gate assembled wild-type (Y526Y) and mutant (Y526F) phages measured in plaque forming units (PFUs) over time (min) graphed on a log10 scale. Purified phage particles were added to a logarithmically growing culture of *E. coli* B at a calculated multiplicity of infection (MOI) of 0.01. The latent period (covering infection, replication, and particle assembly), rise period (particle release and infection of additional cells), and constant period (clearing of bacterial cells resulting in an end of phage propagation) are indicated at the top of the graph. Burst size is calculated from the average PFU in constant period - average PFU in latent period / number of cells for a measure of PFU per infected cell. All values plotted are the average of three technical replicates, and standard deviation is displayed in error bars. Significance according to pairwise t-tests between Y526Y and Y526F with Bonferroni correction is indicated by brackets. Y526L mutant phage did not produce plaques on *E. coli* B and therefore could not be measured in this assay. Created in BioRender. Keown, R. (2026) https://BioRender.com/rpcfqd4

## DISCUSSION

### Implications of PolA motif B mutations for phage biology

Prior analyses of virome data found that a key mutation in motif B of PolA (specifically position 762 [*E. coli*] or 526 [T7]) from unknown virioplankton populations tightly associates with major phylogenetic clades (Schmidt at al., 2014). Subsequent work with a larger number of PolA sequences further substantiated this work (Keown et al., 2022; Nasko et al., 2018) demonstrating that PolA shows greater diversity within viruses than bacteria and that the amino acid identity of position 526/762 dictates broader gene neighbor associations, in particular with helicases and ribonucleotide reductase. These findings broadly indicated the increased selective pressure on PolA within viruses and led to the question addressed in this study of whether 526/762 motif B mutations have significant impacts on phage infection phenotypes.

The substitution of tyrosine with phenylalanine at position 762 is commonly observed in nature. Within large clades of viral sequences, occasional reversions of tyrosine (blue branches) to phenylalanine (red branches) or vice versa can be observed (Fig. 4). These natural variations underscore the evolutionary significance of this amino acid substitution and suggest its potential adaptive value across diverse biological contexts. Conversely, reversion between phenylalanine and leucine are rare, and reversion from tyrosine to leucine has not been observed. Namely, the two leucine clades defined in Keown et al., 2022 (purple branches), are monophyletic and leucine is nearly 100% conserved (Fig. 4). This suggests that efficient enzyme function with leucine at 526/762 requires more compensatory mutations, and once those occur, reversion to other amino acids may not be possible. This need for other compensatory changes within PolA as well as different associated replicon genes, such as helicase (Nasko et al., 2018), are likely the underlying reasons for the *in vivo* failure of the Y526L mutation in the T7 model system. As phylogenetic analysis shows, F526 PolA sequences exist near the base of each leucine clade (Fig. 4). Future work using bacteriophage model systems with wild-type F526 PolA genes and leveraging GGA synthetic biology approaches may be capable of assessing the phenotypic impacts of leucine mutations (i.e. F526L) within phages. This is an important consideration as in various environments, such as the Chesapeake Bay estuary (Schmidt et al., 2014), L526 PolAs were shown to be dominant within the virioplankton.

**Figure 4.**
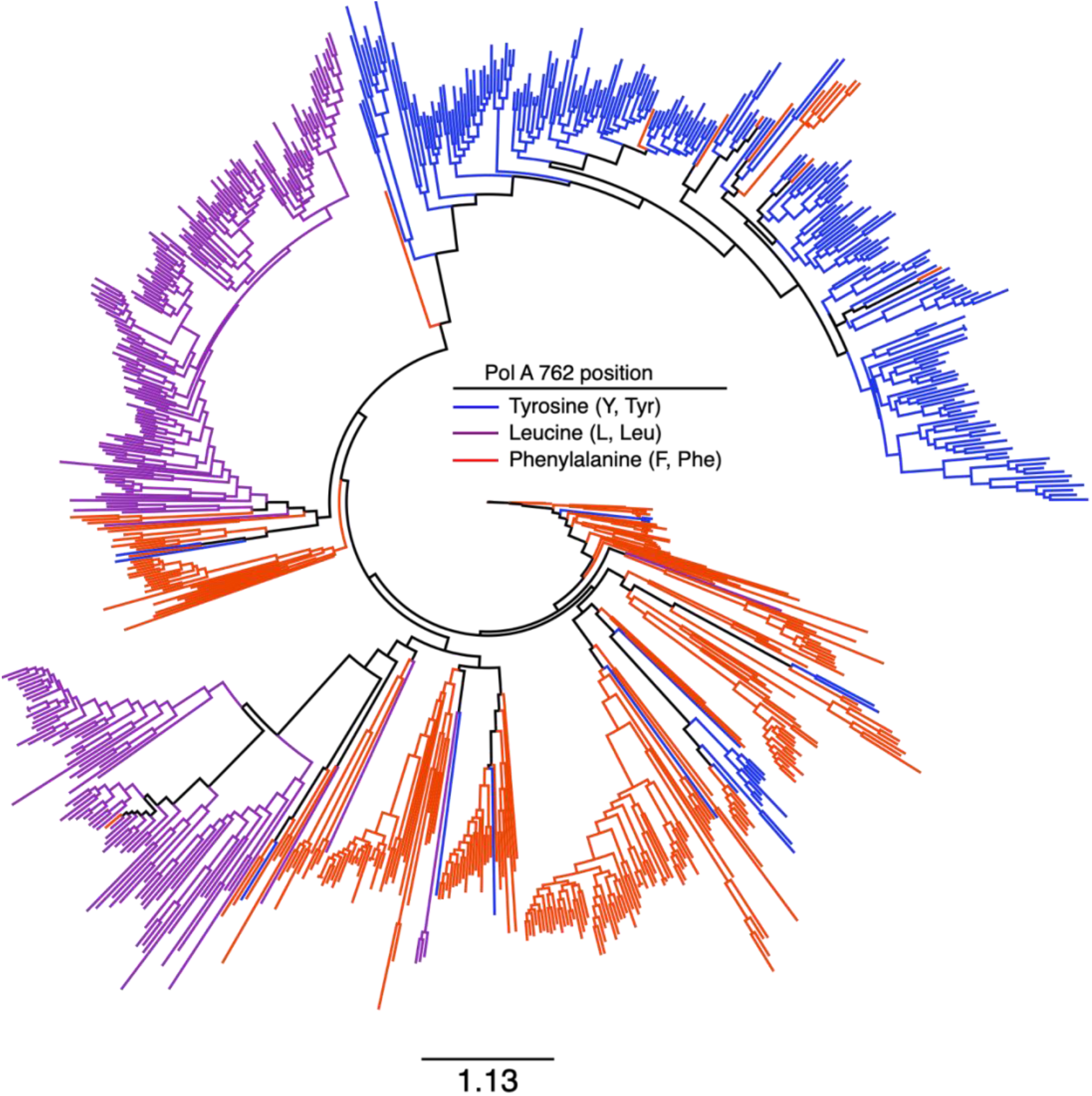
Reversions from tyrosine to phenylalanine are common in nature, while leucine clades are monophyletic. Approximate maximum likelihood tree of 770 PolA amino acid sequences from metagenomic data trimmed to the polymerase domain (DNA_pol_A Superfamily, acc. cl02626). Trimmed sequences were aligned and alignment used to build phylogeny. Branch coloring indicates residue identity at the 762 position (*E. coli* numbering). Scale bar represents the number of amino acid substitutions per site. 762 position identity maps to major phylogenetic clades with leucine maintaining monophyly, while reversions between tyrosine and phenylalanine are common.

### Enzyme biochemistry reflects *in vivo* replication activity

The findings of this study support and extend the seminal work of Tabor and Richardson (1987 and 1995), which explored the *in vitro* biochemical implications of key amino acid residues within the T7 PolA and other DNA polymerases. These studies revealed that the Y526F mutation resulted in decreased discrimination between dNTPs and ddNTPs, underscoring the critical role of the wild-type tyrosine in nucleotide selectivity, and defining T7 DNA polymerase as the best candidate for chain-terminating DNA sequencing (Sanger et al., 1977; Zhu, 2014). Furthermore, these observations hinted at the importance of tyrosine for enzyme nucleotide incorporation efficiency, with the leucine mutant displaying notably lower efficiency. Building upon their foundational *in vitro* work, our data reveal a significant reduction in specific activity for both the phenylalanine and leucine mutants (Fig. 1). Specifically, a 50% decrease in specific activity in dsDNA synthesis was observed for the phenylalanine mutant compared to wild-type tyrosine, suggesting that the hydroxyl on the tyrosine R-group plays a significant role in the rates of dNTP incorporation (Fig. 5). The leucine mutant enzyme exhibited an even more pronounced 97% reduction in specific activity, indicating significant impairment in DNA synthesis efficiency (Fig. 1).

**Figure 5.**
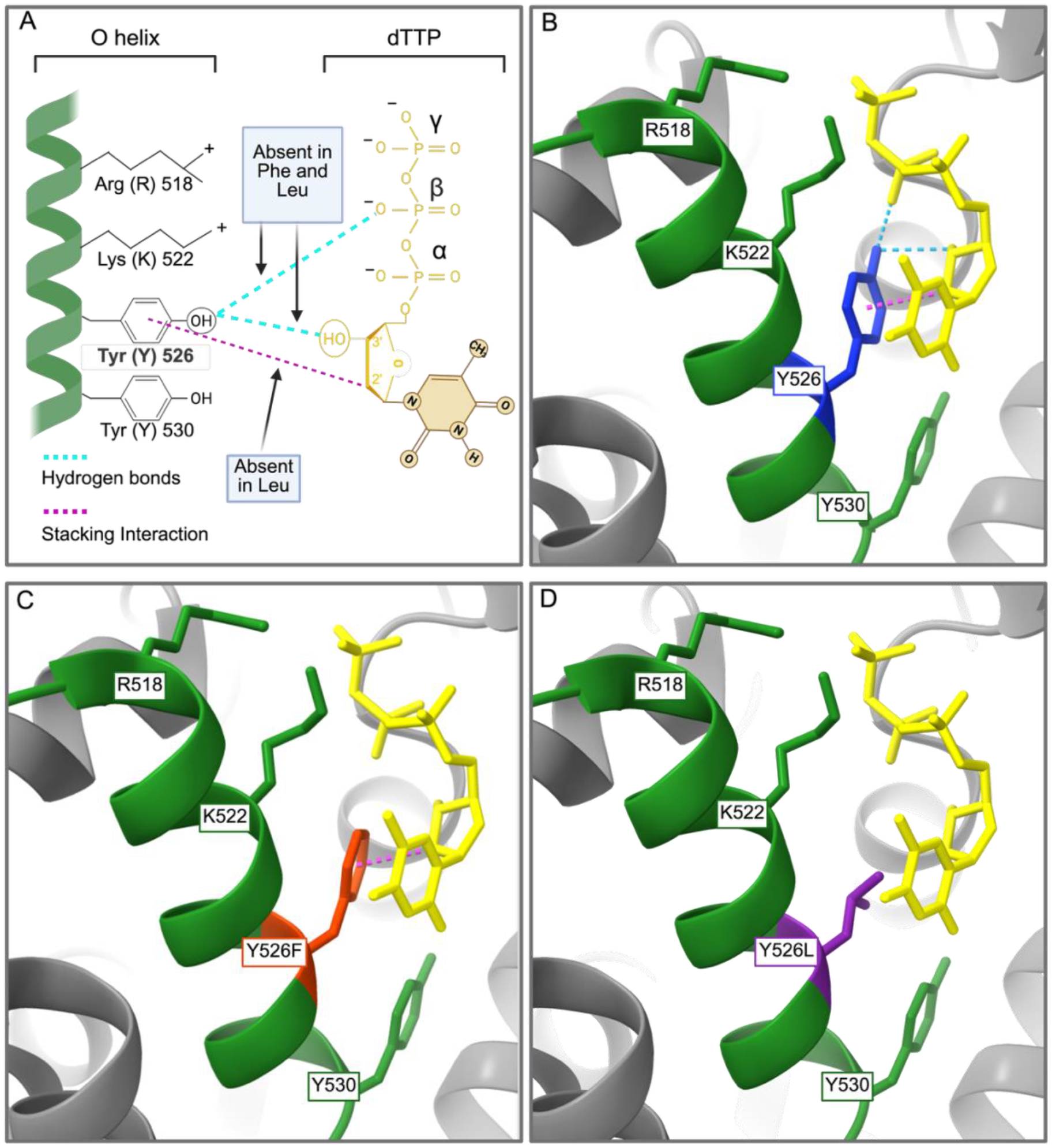
The active site of T7 PolA predicts the loss of molecular interactions between mutant 526 side chains and incoming nucleotides. Interactions between PolA and an incoming dTTP. A) Adapted from Tabor and Richardson, 1995. Modeled interactions are shown between the incoming dNTP and B) Y526, C) Y526F, and D) Y526L. Blue dashed lines represent hydrogen bonding, and purple dashed lines represent interactions with the ribose 2′ carbon. Models show three interactions between wild-type (Y526) and the incoming dTTP, while Y526F loses the ability to H-bond to the dTTP. Y526L additionally loses the ribose interaction, reinforcing that mutation of this residue disrupts the wild-type functionality. Modeled and colored in ChimeraX from PDB ID 6P7E. Created in BioRender. Keown, R. (2026) https://BioRender.com/l24hlhu

Temperature sensitivity of polymerase mutants may provide insights into the environmental adaptability of bacteriophages linked with PolA biochemistry. Previous studies have demonstrated that temperature significantly impacts bacteriophage-mediated lysis efficiency, highlighting the importance of bacterial temperature history (Ameh et al., 2022). The phenylalanine mutant PolA showed a marginal decrease in high temperature tolerance as compared to wild-type, while the leucine mutant enzyme exhibited a drastically limited range of operational temperatures for polymerase activity (Supplementary Fig. S5). *E. coli* can tolerate temperatures up to 45 °C (Kumar and Libchaber, 2013), and the data suggest that T7 phages with wild-type PolA would likely withstand high temperature stress more effectively than strains with phenylalanine at this position. The leucine mutant enzyme did not tolerate any temperature outside the *E. coli* 37°C optimum, further confirming the low specific activity of this mutant PolA (Fig. 1 and Supplementary Fig. S5). It is possible that within nature, phages carrying Y526 PolA genes demonstrate greater temperature adaptability than those with either F526 or L526 PolAs. However, the predominance of L526 PolAs within Chesapeake virioplankton (Schmidt et al., 2014) counters this idea as the Chesapeake experiences substantial seasonal variability in water temperature.

Surprisingly, no differences in enzyme fidelity were observed when comparing the wild-type and mutant phenylalanine enzymes (data unavailable from the Y526L mutant due to low polymerase efficiency) (Supplementary Figs. S6 and S7). An earlier study observed that Y526 and F526 PolAs predominantly occurred within virulent phages and that L526 PolAs occurred within temperate phages (Schmidt et al., 2014). This led to the hypothesis, supported by earlier work using a *Thermus aquaticus* model (Suzuki et al., 2000), that within phages there exists a trade-off between polymerase nucleotide incorporation efficiency and fidelity. Along this hypothesized spectrum, tyrosine PolAs would have the highest efficiency and the lowest fidelity and leucine PolAs would have the lowest efficiency and highest fidelity, with phenylalanine PolAs existing between these extremes. However, fidelity assays did not support the existence of an efficiency/fidelity trade-off between tyrosine and phenylalanine PolA mutants *in vitro*. It is still possible that this trade-off exists *in vivo* and alternate bacteriophage model systems with different suites of accessory replication genes could provide the experimental means for testing this hypothesis.

The *in vitro* experiments provided context essential for understanding how mutations that affect enzyme function could translate into changes in viral fitness and adaptability, which are important factors in the evolutionary dynamics of phage populations in nature (Díaz-Muñoz et al., 2014; De Leeuw et al., 2020). *In vitro* data revealed striking parallels with *in vivo* findings. Specifically, the phenylalanine 526 mutation exhibited a 50% reduction in enzymatic activity and displayed a corresponding 50% decrease in burst size (i.e., phage production) and a significant increase in latent period compared with the Y526 wild-type (Fig. 3). In addition, the 97% reduction in polymerase activity observed in the L526 enzyme (Fig. 1) reflects the total lack of phage production *in vivo* (Fig. 2). Quantitative PCR confirms that *in vivo* DNA replication by the L526 variant is negligible; consequently, *E. coli* cell division likely outpaces viral replication, resulting in non-productive infection. This convergence of results across *in vitro* and *in vivo* assays strengthens the validity of the findings and underscores the critical role of residue 526 in determining both enzyme function and viral life history.

### Impact of position 526 on nucleotide selectivity and incorporation

Previous biochemical studies of PolAs have identified site-specific amino acids that limit the incorporation of ribonucleotides (rNTPs) i.e., the “steric gate” (Astake et al., 1998). In *E. coli* Pol I, the steric gate is Glu710, and alignment of the T7 sequence shows Glu480 to be conserved (data not shown). The bulky aromatic ring of phenylalanine at position 762 in *E. coli* Pol I was determined to be critical for constraining the incoming nucleotide, acting as a steric gate assistant. Specifically, the phenylalanine (and by extension, tyrosine) aromatic ring interacts with the 2′ carbon of the incoming nucleotide to position the ribose for interrogation by the Glu480 steric gate while helping to position the nucleotide for catalysis. Loss of this function through the substitution of a non-bulky, non-aromatic residue such as leucine at 762/526 likely impairs proper positioning of the incoming nucleotide and leads to the observed decrease in polymerization efficiency observed in the leucine mutant. The leucine substitution could also lead to decreased fidelity through the disruption of the steric gate and further work is needed to determine whether leucine-containing polymerases incorporate rNTPs.

As shown in earlier work (Tabor and Richardson, 1995), the hydroxyl group on the tyrosine residue is critical for distinguishing between ddNTPs and dNTPs. Substitution with phenylalanine results in a loss of these interactions, notably the loss of hydrogen bonding to the beta phosphate, but it maintains the interactions through the aromatic ring (Fig. 5). The Y526L mutant lacks an aromatic system and lacks significant interactions with the dNTP substrate. These findings suggest that for native leucine 526 polymerases, as observed in viral metagenomes (Schmidt et al., 2014; Nasko et al., 2018; Keown et al., 2022), compensatory mutations are necessary to fulfill the role of Y526 in nucleotide positions and steric gate interrogation.

### Applications of Golden Gate Assembly in phage genome engineering

Leveraging the High Complexity Golden Gate Assembly tools (Sikkema et al., 2023), this study successfully produced and characterized mutant T7 phages. While the catalytic position 526 has long been of interest for biotechnology and enzymology (Tabor and Richardson, 1987 and 1995; Suzuki et al., 2000), this study addressed the broader genotype-to-phenotype question of how 526 PolA mutations might impact phage infection dynamics. Previously, incorporating single amino acid changes into the T7 genome via recombineering was cost and labor intensive, however, HC-GGA provided a cost-effective means for introducing single codon changes within the context of an entire phage genome. GGA is also capable of larger changes such as gene deletion or substitution, enabling production of mutant phages on a timeline of days to weeks, greatly accelerating the study of bacteriophage biology. Construction of mutant T7 genomes via HC-GGA enabled *in vivo* experiments with the mutations in their full genomic context, providing insights into the impact on viral replication and production within a host environment. This context is essential for understanding how mutations that affect enzyme function translate into changes in viral fitness and adaptability, important factors in the evolutionary dynamics of phage populations in nature (Díaz-Muñoz et al., 2014; De Leeuw et al., 2020).

### Mutation from tyrosine to phenylalanine is common in nature

Correlating this study’s findings with metagenomic data provides insights on the prevalence and evolutionary dynamics of tyrosine-to-phenylalanine mutations within natural viral communities. Although the phenylalanine mutant phage exhibited a reduced burst size (Fig. 3), which might seem disadvantageous, this could potentially confer an evolutionary advantage in environments where host growth rates are slower. The Y526F phage qPCR data indicated a lower gene copy number 30 min post-infection while the wild-type phage was nearly 2 logs higher (Supplementary Fig. S9A). However, it’s plausible that the phenylalanine phage could yield higher final copy number after 2 h, as the extended culture time may provide more opportunities for reinfection after the initial burst around 30 min (Fig. 3 and Supplementary Fig. S9A). Thus, while the phenylalanine mutation initially reduced replication efficiency and burst size, it may ultimately have resulted in a greater number of progeny over time, potentially enhancing the overall fitness of the bacteriophage.

### Implications for bacteriophage synthetic biology studies

Data presented in this study provide strong support for the 526/762 hypotheses described in the introduction and historical literature (Schmidt et al., 2014; Nasko et al., 2018; Keown et al., 2022). By providing detailed insights into how mutations at this position affects DNA synthesis and viral infection dynamics, this work offers a foundation for further exploration of these hypotheses. Investigating compensatory mechanisms in leucine 526/762 PolAs that restore the observed impairments in PolA activity will be a key direction for future research. Identifying additional changes analogous to those seen in naturally occurring leucine PolAs could elucidate how these mutations are accommodated within viral systems. Moreover, exploring the effects of whole gene substitution within T7 bacteriophage, as well as mutations in associated enzymes like helicases, could provide insights to the interactions between different components of the T7 replicon. This approach could enhance our understanding of how replication gene modules contribute to phage fitness and evolution. The T7 system’s versatility and the advances in Golden Gate phage engineering offer promising avenues for these explorations, enabling detailed studies of genetic diversity and its effects on phage biology.

## SUPPLEMENTARY DATA

Supplementary Data are available online.

## AUTHOR CONTRIBUTIONS

Rachel Keown: Conceptualization, Formal analysis, Investigation, Methodology, Validation, Visualization, Writing – original draft, review & editing. Andrew Sikkema: Conceptualization, Investigation, Visualization, Writing—review & editing. Victoria Barbone: Investigation. Barbra Ferrell: Conceptualization, Investigation, Writing— review & editing. Owen Donnelly: Investigation. Sydney Iredell: Investigation. Kelly Zatopek: Investigation, Resources, Writing—review & editing. Phillip Brumm: Investigation, Resources. David Mead: Conceptualization, Supervision, Writing— review & editing. Gregory Lohman: Conceptualization, Supervision, Writing—review & editing. K. Eric Wommack: Conceptualization, Funding acquisition, Supervision, Writing—review & editing. Shawn Polson: Conceptualization, Funding acquisition, Supervision, Writing—review & editing.

## ACKNOWLEDGEMENTS

The authors would like to thank Jacob Dums at the University of Delaware for his contributions to the initial conceptualization of this work. We would also like to thank Spencer Toth at the University of Delaware for her assistance with one-step growth curve data collection. We are grateful to Mark Shaw and the University of Delaware DNA Sequencing and Genotyping Center [RRID:SCR_012230] for their expertise and assistance with library preparation and sequencing. We also extend our gratitude to Kurt Throckmorton, Joyanne MacDonald, Alyssa Hassinger, and Aaron Lomax from Varizymes (Middleton, WI) for their invaluable training in many of the techniques utilized in this study. Thank you to the University of Delaware Bioinformatics Data Science Core [RRID:SCR_017696], Delaware INBRE, and the Delaware Biotechnology Institute for access to computational infrastructure, including the Biomix high performance computational cluster and the BioStoRe data storage resource.

## REFERENCES

Ameh, E. M., Nocker, A., Tyrrel, S., Harris, J. A., Orlova, E. V., & Ignatiou, A. Effect of temperature on bacteriophage-mediated lysis efficiency with a special emphasis on bacterial temperature history. J. Mater. Environ. Sci., 13(9) (2022) 1056–1066

Astatke, M., Ng, K., Grindley, N. D. F., & Joyce, C. M. A single side chain prevents escherichia coli DNA polymerase I (klenow fragment) from incorporating ribonucleotides.

Betancurt-Anzola, L., O’Connell, K. C., Potapov, V., Ong, J. L., Tanner, N. A., Sauguet, L., & Zatopek, K. M. (2025). The A, B, C, D’s of replicative DNA polymerase fidelity: utilizing high-throughput single-molecule sequencing to understand the molecular basis for DNA polymerase accuracy. Nucleic acids research, 53(21), gkaf1143. doi: 10.1093/nar/gkaf1143

Brown, J. A., & Suo, Z. (2011). Unlocking the sugar “Steric gate” of DNA polymerases. Biochemistry, 50(7), 1135. doi:10.1021/bi101915z

Chiu, J., Tillett, D., & March, P. E. (2005). Coexpression of the subunits of T7 DNA polymerase from an artificial operon allows one-step purification of active gp5/trx complex. Protein Expression and Purification, 47(1), 264. doi:10.1016/j.pep.2005.10.016

De Leeuw, M., Baron, M., Ben David, O., & Kushmaro, A. (2020). Molecular insights into bacteriophage evolution toward its host MDPI AG. doi:10.3390/v12101132

Díaz-Muñoz, S. L., & Koskella, B. (2014). Bacteria–Phage interactions in natural environments Elsevier. doi:10.1016/b978-0-12-800259-9.00004-4

Keown, R. A., Dums, J. T., Brumm, P. J., Macdonald, J., Mead, D. A., Ferrell, B. D., Moore, R. M., Harrison, A. O., Polson, S. W., and Wommack, K. E. (2022). Novel viral DNA polymerases from metagenomes suggest genomic sources of strand-displacing biochemical phenotypes Frontiers Media SA. doi:10.3389/fmicb.2022.858366

Kropinski, A. M. (2018). Practical advice on the one-step growth curve. Bacteriophages (pp. 41–47). New York, NY: Springer New York. doi:10.1007/978-1-4939-7343-9_3 Retrieved from http://link.springer.com/10.1007/978-1-4939-7343-9_3

Kumar, P., & Libchaber, A. (2013). Pressure and temperature dependence of growth and morphology of escherichia coli: Experiments and stochastic model. Biophysical Journal, 105(3), 783. doi: 10.1016/j.bpj.2013.06.029

Lee, S., & Richardson, C. C. (2012). Choreography of bacteriophage T7 DNA replication. Current Opinion in Chemical Biology, 15(5), 580. doi:10.1016/j.cbpa.2011.07.024

Lennox, E. S. (1955). Transduction of linked genetic characters of the host by bacteriophage P1. Virology, 1(2), 190–206. doi:10.1016/0042-6822(55)90016-7

Makiela-Dzbenska, K., Jaszczur, M., Banach-Orlowska, M., Jonczyk, P., Schaaper, R. M., & Fijaklowska, I. J., (2009). Role of *Escherichia coli* DNA polymerase I in chromosomal DNA replication fidelity. Mol. Microbiol. 74, 114–1127. doi: 10.1111/j.1365-2958.2009.06921.x

Mark, D. F., & Richardson, C. C. (1976). Escherichia coli thioredoxin: a subunit of bacteriophage T7 DNA polymerase. Proceedings of the National Academy of Sciences of the United States of America, 73(3), 780–784. doi: 10.1073/pnas.73.3.780

Moore, R. M., Harrison, A. O., Mcallister, S. M., Polson, S. W., & Wommack, K. E. (2020). Iroki: Automatic customization and visualization of phylogenetic trees PeerJ. doi:10.7717/peerj.8584

Nasko, D. J., Chopyk, J., Sakowski, E. G., Ferrell, B. D., Polson, S. W., & Wommack, K. E. (2018). Family A DNA polymerase phylogeny uncovers diversity and replication gene organization in the virioplankton. Frontiers in Microbiology, 9 doi:10.3389/fmicb.2018.03053

Ong, S., Ghode, P., Narenderan, A., Lao, S., Willenborg, F., Eden, T. V., Marsh, C. O., Yew, W. S., & Fredens, J. (2026). Bridging continuous and discrete evolution through a controllable, hypermutagenic phage-bacteria system. Nature Microbiology, 1–13. doi: 10.1038/s41564-026-02346-y

Parker, D. R., Sikkema, A. P., Anderson, R. K., Lohman, G. J. S., & Nugen, S. R. (2025). Recoded bacteriophage genome for bio-orthogonal-enabled concentration and detection of *E. coli* in drinking water. ACS Synthetic Biology, 15(1), 233–242. doi: 10.1021/acssynbio.5c00665

Potapov, V., Fu, X., Dai, N., Corrêa, I. R., Tanner, N. A., & Ong, J. L. (2018). Base modifications affecting RNA polymerase and reverse transcriptase fidelity. Nucleic Acids Research, 46(11), 5753. doi:10.1093/nar/gky341

Pryor, J. M., Potapov, V., Bilotti, K., Pokhrel, N., & Lohman, G. J. S. (2022). Rapid 40 kb genome construction from 52 parts through data-optimized assembly design. ACS Synthetic Biology, 11(6), 2036. doi:10.1021/acssynbio.1c00525

Pryor, J. M., Potapov, V., Kucera, R. B., Bilotti, K., Cantor, E. J., & Lohman, G. J. S. (2020). Enabling one-pot golden gate assemblies of unprecedented complexity using data-optimized assembly design. Plos One, 15(9) doi:10.1371/journal.pone.0238592

Richardson, C. C. (2015). It seems like only yesterday. Annual Review of Biochemistry, 84(1), 1. doi:10.1146/annurev-biochem-060614-033850

Rosario, K., & Breitbart, M. (2011). Exploring the viral world through metagenomics. Current Opinion in Virology, 1(4), 289–297. 10.1016/j.coviro.2011.06.004

Roux, S., Matthijnssens, J., & Dutilh, B. E. (2021). Metagenomics in Virology. Encyclopedia of Virology, 133–140. 10.1016/B978-0-12-809633-8.20957-6

Sanger, F., Nicklen, S., & Coulson, A. R. (1977). DNA sequencing with chain-terminating inhibitors. Proceedings of the National Academy of Sciences - PNAS, 74(12), 5463–5467. doi:10.1073/pnas.74.12.5463

Schmidt, H. F., Sakowski, E. G., Williamson, S. J., Polson, S. W., & Wommack, K. E. (2014). Shotgun metagenomics indicates novel family A DNA polymerases predominate within marine virioplankton. The ISME Journal, 8(1), 103. doi:10.1038/ismej.2013.124

Sikkema, A. P., Tabatabaei, S. K., Lee, Y., Lund, S., & Lohman, G. J. S. (2023). High-Complexity One-Pot golden gate assembly. Current Protocols, 3(9) doi:10.1002/cpz1.882

Suttle, C. A. (2005). Viruses in the sea. Nature, 437(7057), 356–361. doi:10.1038/nature04160

Suttle, C. A. (2007). Marine viruses — major players in the global ecosystem. Nature Reviews Microbiology, 5(10), 801–812. doi:10.1038/nrmicro1750

Suzuki, M., Yoshida, S., Adman, E. T., Blank, A., & Loeb, L. A. (2000). Thermus aquaticus DNA polymerase I mutants with altered fidelity. Journal of Biological Chemistry, 275(42), 32728. doi:10.1074/jbc.m000097200

Tabor, S., & Richardson, C. C. (1995). A single residue in DNA polymerases of the escherichia coli DNA polymerase I family is critical for distinguishing between deoxy- and dideoxyribonucleotides. Proceedings of the National Academy of Sciences, 92(14), 6339–6343. doi:10.1073/pnas.92.14.6339

Tabor, S., & Richardson, C. C. (1987). DNA sequence analysis with a modified bacteriophage T7 DNA polymerase. Proceedings of the National Academy of Sciences - PNAS, 84(14), 4767–4771. doi:10.1073/pnas.84.14.4767

Zhu, B., Schnorr, K. M., Hamdan, S., & Abdullah, K. (2014). Bacteriophage T7 DNA polymerase – sequenase. Frontiers in Microbiology, 5 doi:10.3389/fmicb.2014.00181

