## Supplementary Materials for "Single amino acid substitution in DNA Polymerase I dramatically alters infection dynamics of bacteriophage T7"

### Supplementary Figures

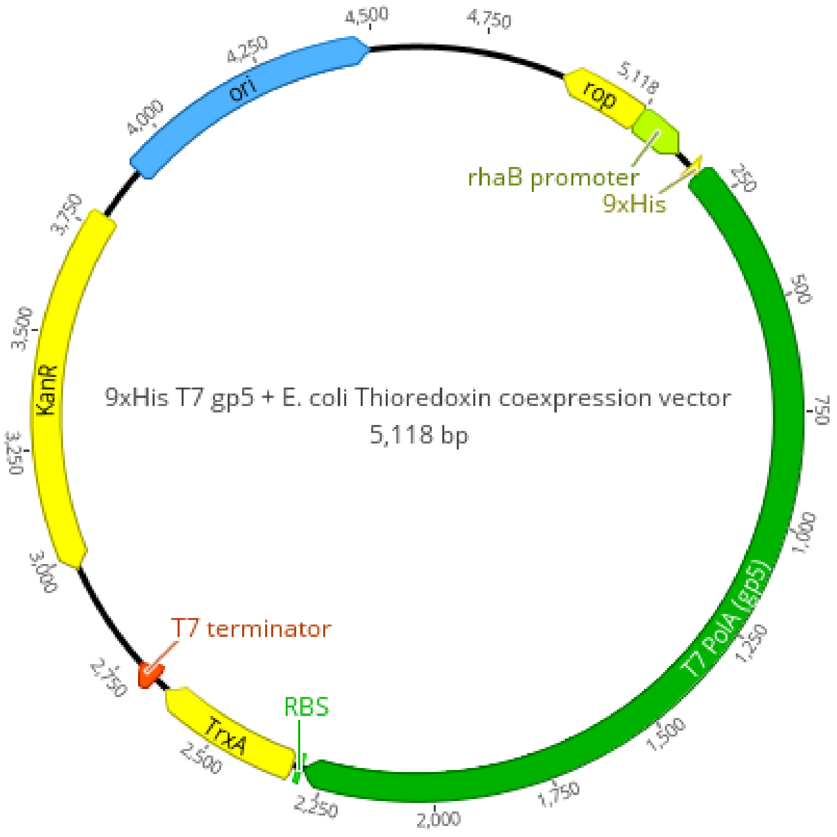


Supplementary Figure S1. Plasmid map for 9xHis-tagged T7 PolA and thioredoxin co-expression

The T7 DNA polymerase gene (gp5) was flanked with a 9x Histidine tag at the N-terminus and the *E. coli* thioredoxin gene (*trxA*) at the C-terminus, and inserted into a custom rhamnose-inducible backbone. Sequence was annotated and visualized in Geneious v2024.0.5 (https://www.geneious.com).

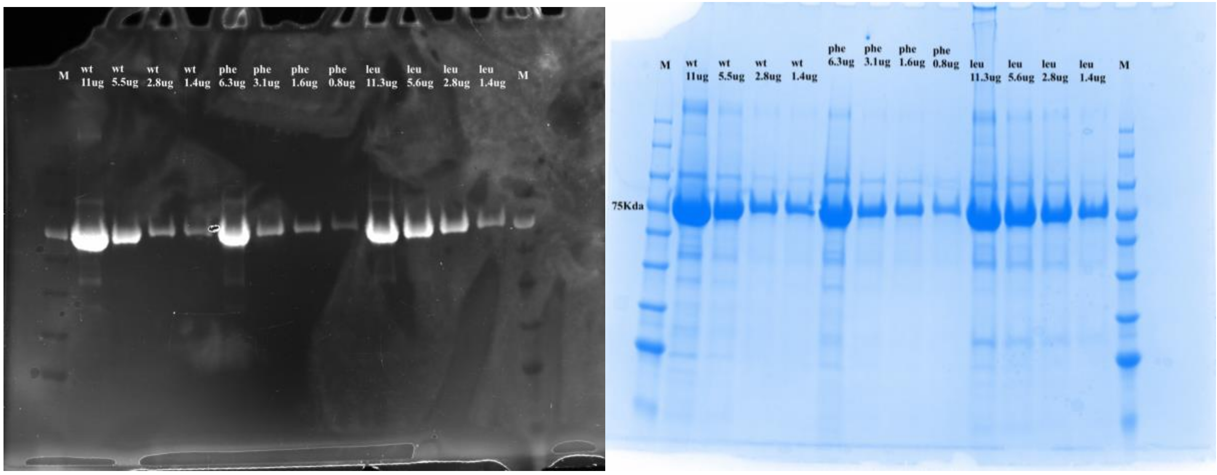


Supplementary Figure S2. SDS-PAGE gel of T7 PolA–TrxA protein stocks stained with His-Stain (ThermoFisher Scientific) and Coomassie blue (Research Products International), respectively.

Cultures were grown for 18 h at 30 °C, 200 rpm in 100 mL LB, 30 µg/mL kanamycin, 0.4% rhamnose (in place of dextrose). Cultures were harvested by centrifugation, lysed by sonication, and analyzed by SDS-PAGE on a 4–20% gradient gel. Expected molecular weight ~75 kDa.

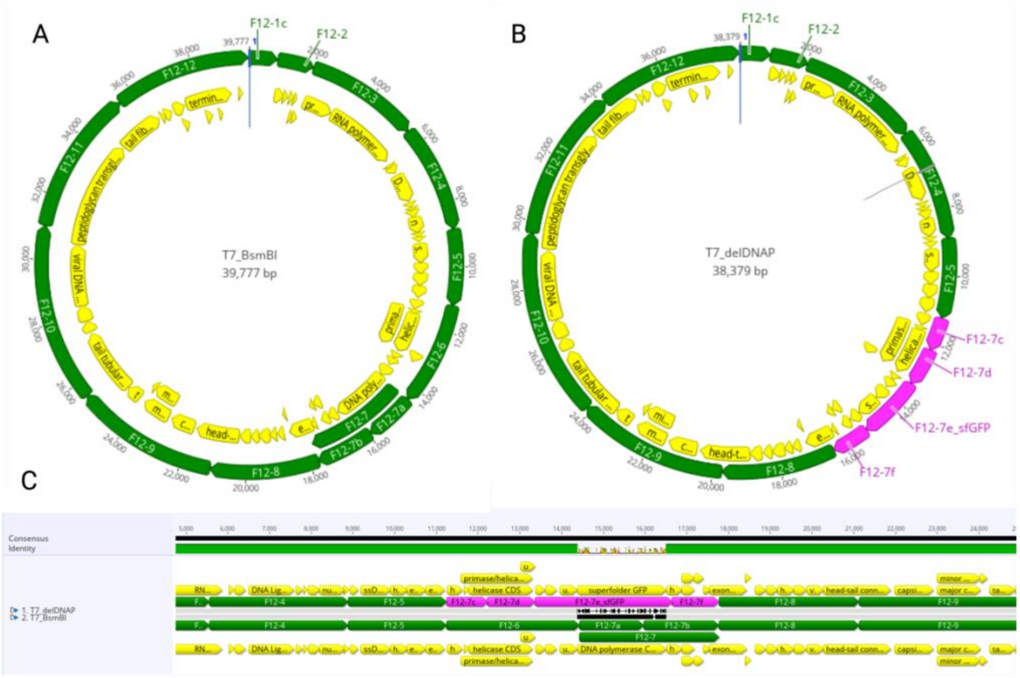


Supplementary Figure S3. Genome maps of domesticated T7 and ∆DNAP, and alignment of assemblies

Genome maps of A) Domesticated T7 genome with 13 fragments (green) and B) ∆DNAP genome with 10 original fragments and 4 new fragments (pink). C) Alignment of domesticated and ∆DNAP assemblies where GFP was substituted for DNA polymerase, maintaining start and stop codons. Sequences were annotated and visualized in Geneious v2024.0.5 (https://www.geneious.com).

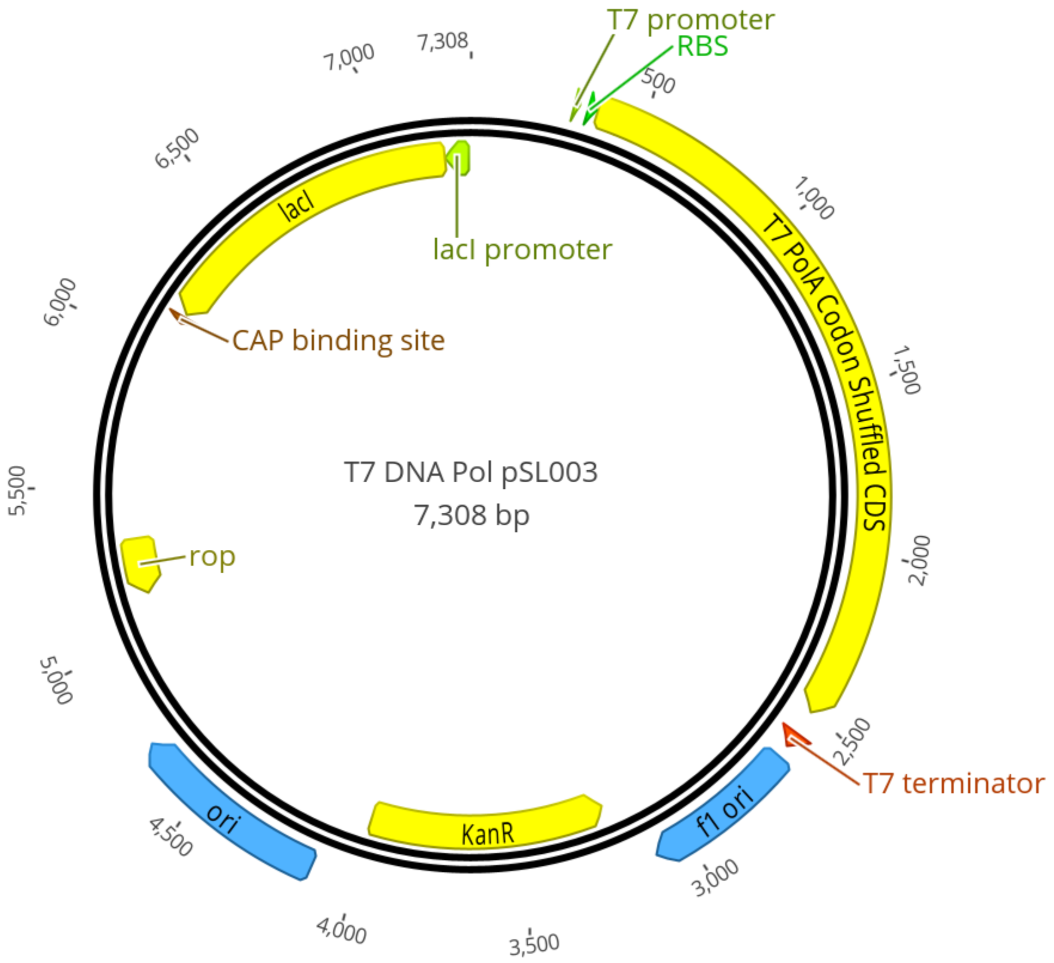


Supplementary Figure S4. Plasmid map of rescue vector

The T7 DNA polymerase gene (gp5) was inserted untagged into a custom pET backbone that lacked the LacO site. Sequence was annotated and visualized in Geneious v2024.0.5 (https://www.geneious.com).

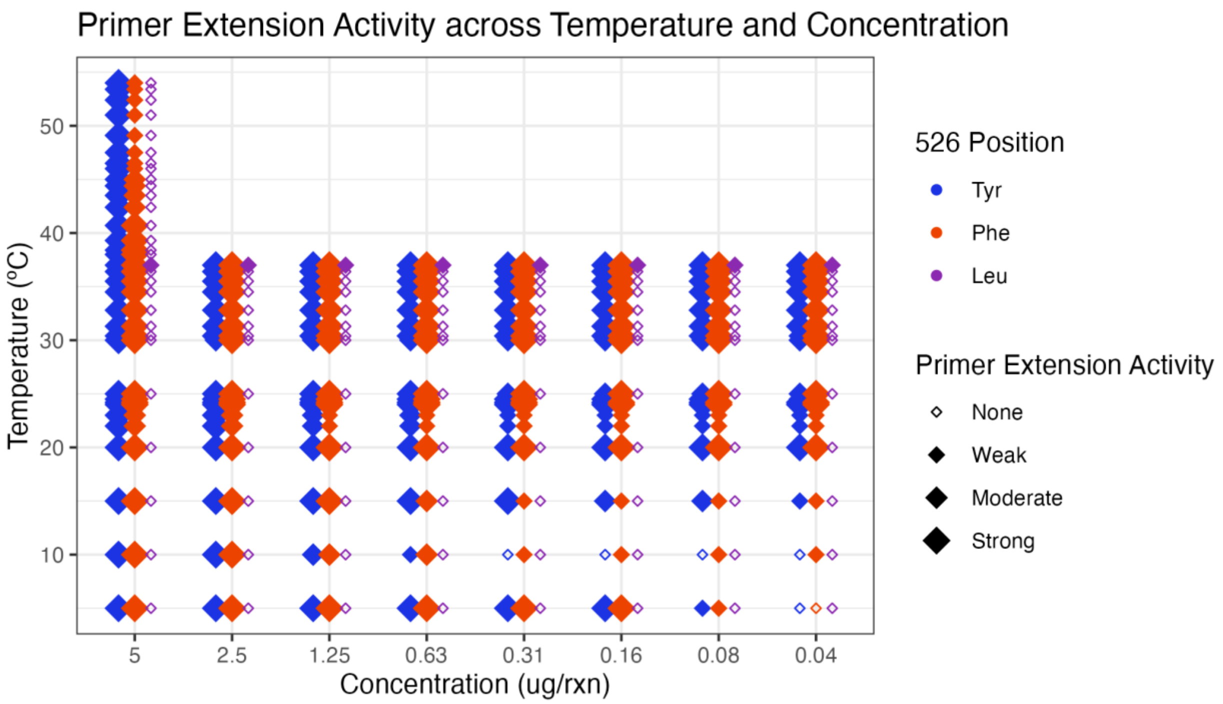


Supplementary Figure S5. Substitution of amino acid 526 alters temperature tolerance in T7 DNA polymerase.

Primer extension (PE) activity for all mutant T7 DNA polymerases graphed over temperature (y-axis) and concentration (x-axis) gradients. Positive products were qualitatively placed into categories of strong, moderate, or weak activity. Mutant enzyme Y526F showed slightly decreased activity in the high temperature range (>45 °C). Mutant enzyme Y526L showed minimal activity outside of the enzyme optimum (37 °C).

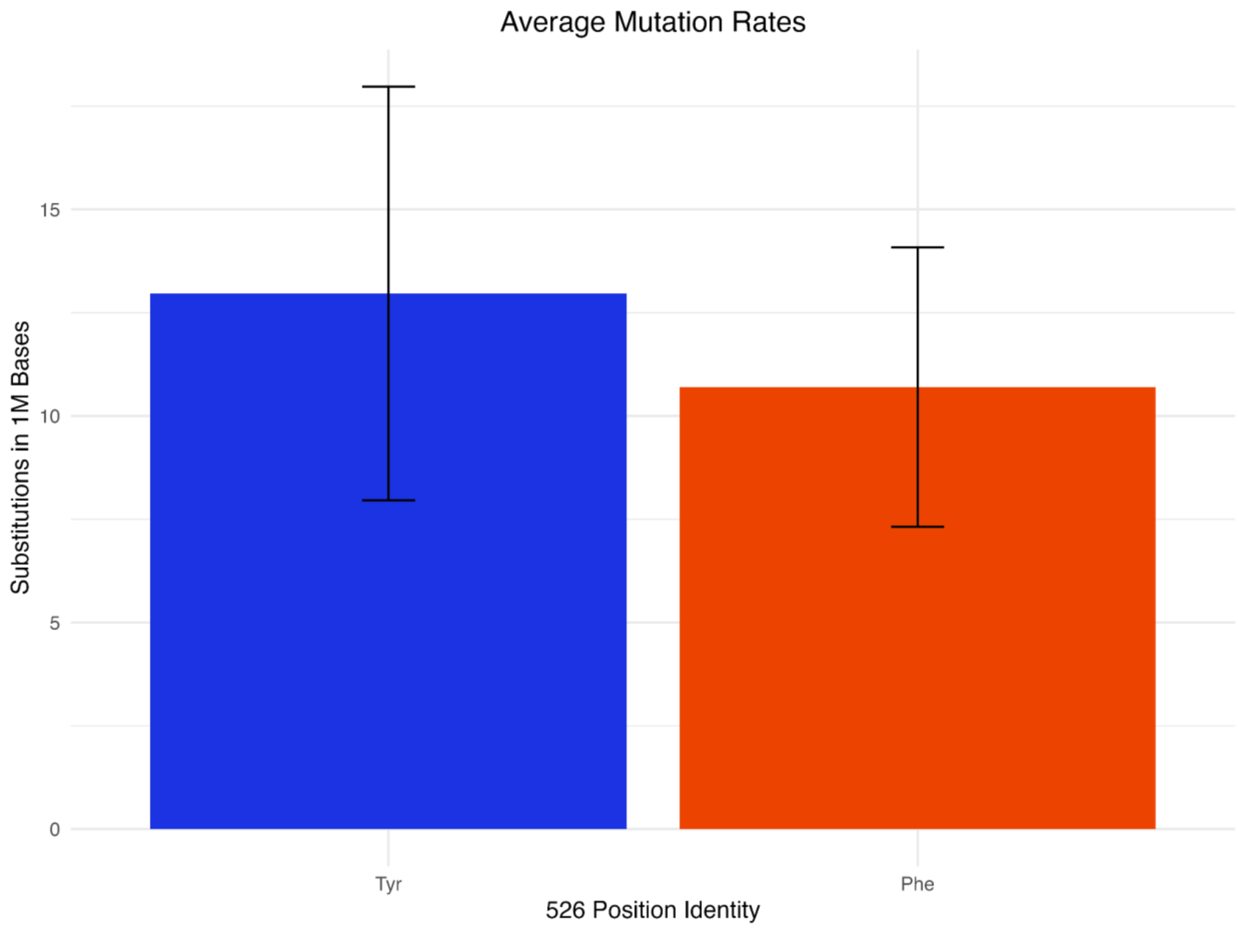


Supplementary Figure S6. Substitution rate per 1M bases

Mutation rates for wild-type (Y526Y) and mutant (Y526F) T7 DNA polymerase proteins were measured with a PacBio Sequencing assay. Final values were calculated in substitutions per 1M bases. The averages of five replicates are reported with error bars denoting one standard deviation.

Supplementary Figure S7. Substitution Spectra for T7 DNA polymerases

Rates of substitution type, reported here as a percentage of 100.

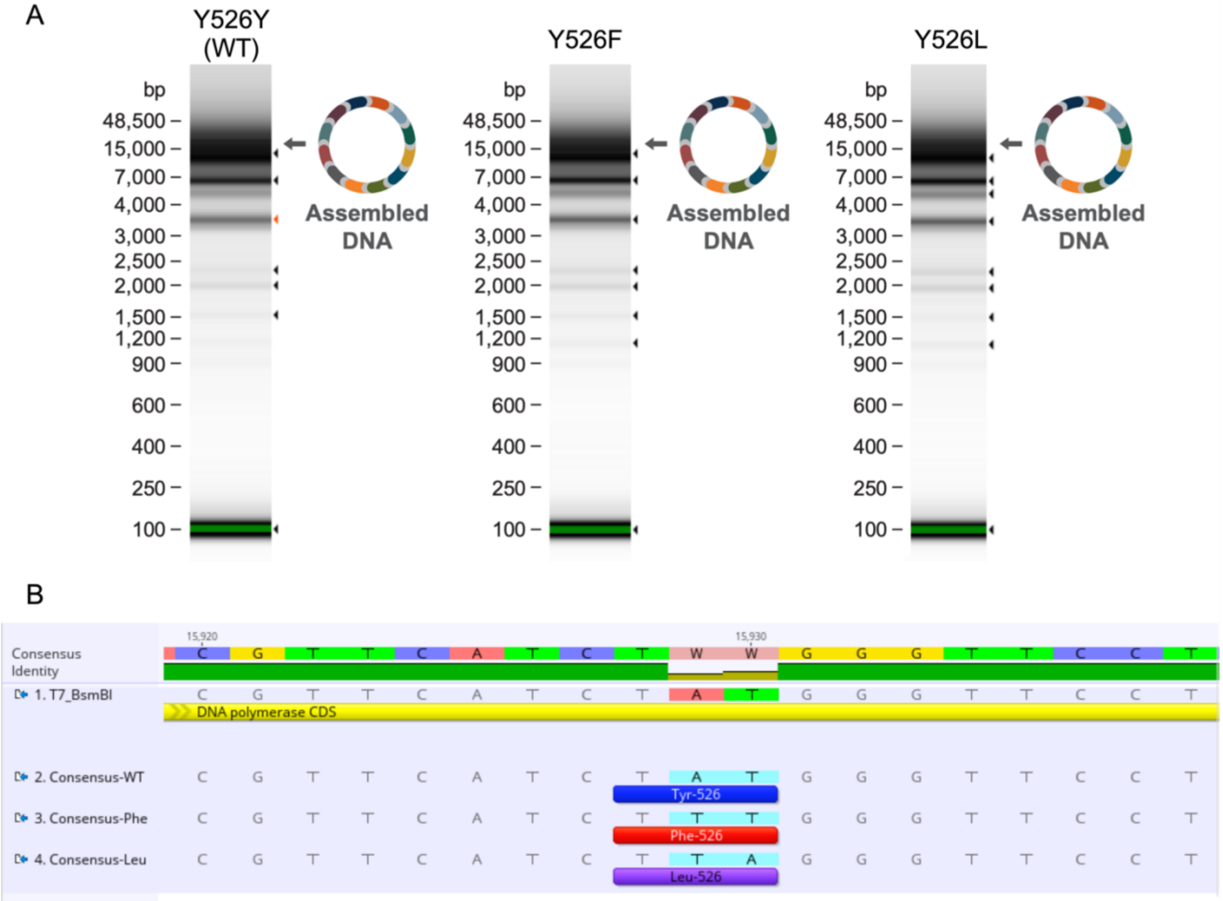


Supplementary Figure S8. Capillary electrophoresis and Illumina sequencing of Golden Gate assembled phage genomes

A) Capillary electrophoresis traces (Agilent TapeStation, Santa Clara, CA) of completed assemblies. B) Consensus sequences assembled from Illumina 2x151 bp reads mapped to and aligned with reference genome (domesticated T7). Codon for amino acid 526 is translated, annotated, and visualized in Geneious v2024.0.5 (https://www.geneious.com).

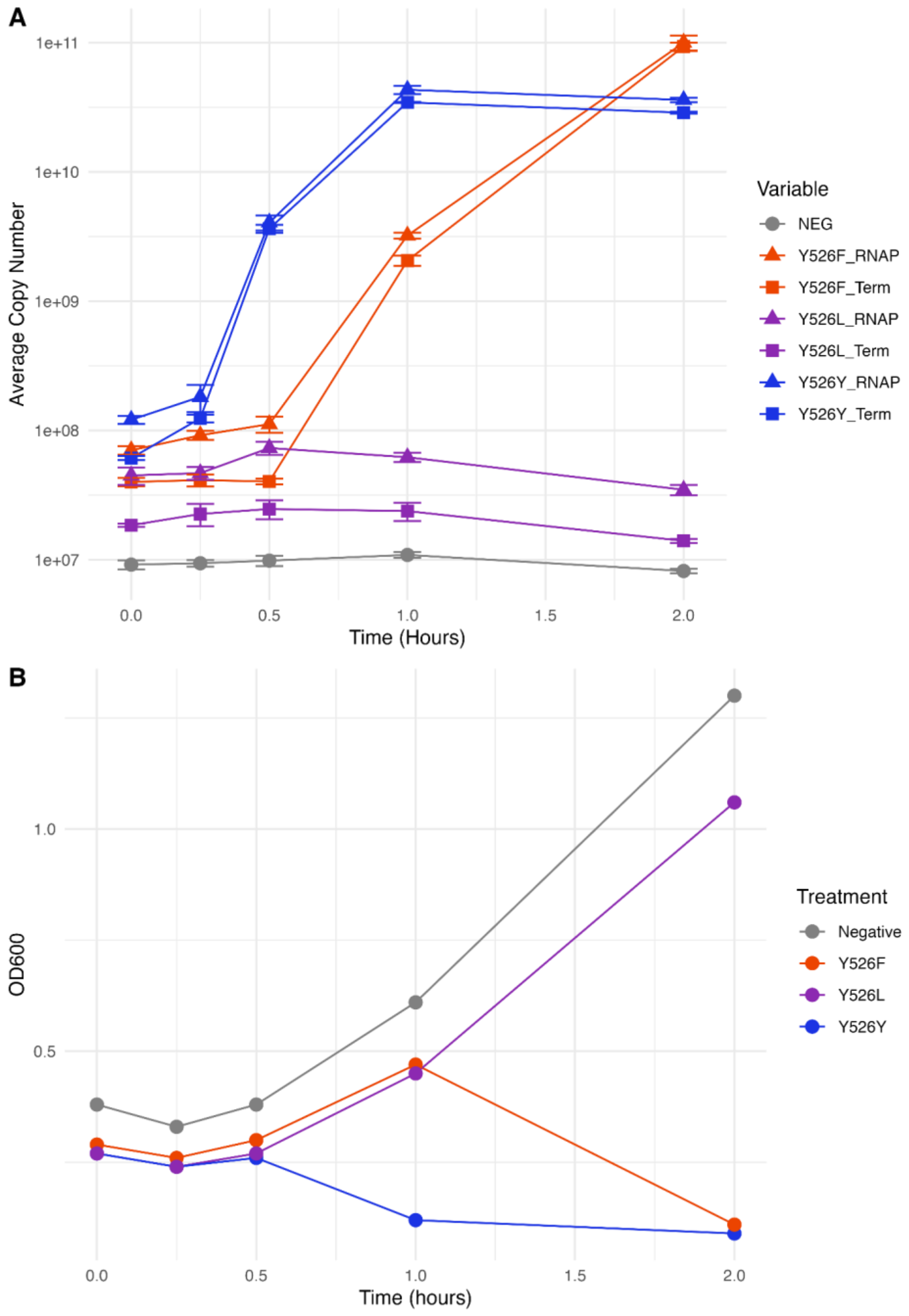


Supplementary Figure S9. Genome replication and cell lysis phenotypes change for PolA mutant phage.

A) Gene copy number determined by qPCR for two T7 genomic targets during infection: RNA polymerase (RNAP) and terminase large subunit (Term). B) Optical density (600 nm) of each culture. Cells were transformed by electroporation, recovered for 90 minutes, and T0 was marked when outgrowth was inoculated into equilibrated LB broth. Cultures were incubated at 37 °C for 32 h and sampled at regular intervals. Data was truncated to the first 2 h (five timepoints).

### Supplementary Tables

#### Supplementary Table S1. Primer sequences

| Primer Name | Primer Sequence (5’->3’)* | Application |
|---|---|---|
| 39751F | ggctaccgtctcgAATATACCATAAAAATCTGAGTGACTAT C | Phage Genome Fragment Generation |
| 874R | ggctaccgtctctGATACCCTTGAGTTATCCGC | Phage Genome Fragment Generation |
| 871F | ggctaccgtctctTATCGCAAGGTGCCCTTTATG | Phage Genome Fragment Generation |
| 1998R | ggctaccgtctctTTATGAGATAGCGTTCAGTGTGTTG | Phage Genome Fragment Generation |
| 1995F | ggctaccgtctctATAACGAACATAAAGGACACAATGC | Phage Genome Fragment Generation |
| 5534R | ggctaccgtctccACCGTCTTGGCTGTGTAC | Phage Genome Fragment Generation |
| 5531F | ggctaccgtctcaCGGTAGCCACCTTCGTAAG | Phage Genome Fragment Generation |
| 8857R | ggctaccgtctctTGTACGCATCGTCGAATGTCTC | Phage Genome Fragment Generation |
| 8854F | ggctaccgtctccTACATCAAGCTGTATGAACTTTGG | Phage Genome Fragment Generation |
| 11208R | ggctaccgtctcgCGAACAAAGGGAAACCGCTGTG | Phage Genome Fragment Generation |
| 11205F | ggctaccgtctctTTCGCATTGGAGGTCAAATAATGC | Phage Genome Fragment Generation |
| 14405R | ggctaccgtctccAACTTAGTGACGCTCTCTAAGAG | Phage Genome Fragment Generation |
| 14402F | ggctaccgtctccAGTTCCACTGCGGGGTTATC | Phage Genome Fragment Generation |
| 15940R | ggctaccgtctccAGAGGAACCCATAGATGAACGTC | Phage Genome Fragment Generation |
| 15940R-A15929T** | ggctaccgtctccAGAGGAACCCAAAGATGAACGTCTTAGC | Phage Genome Fragment Generation |
| 15940R-A15929T-T15930A** | ggctaccgtctccAGAGGAACCCTAAGATGAACGTCTTAGC | Phage Genome Fragment Generation |
| 15937F | ggctaccgtctcaCTCTATGGTGCTGGTGATGAGAAG | Phage Genome Fragment Generation |
| 17736R | ggctaccgtctccGCACCTGCCCAAGCCTTC | Phage Genome Fragment Generation |
| 17733F | ggctaccgtctccGTGCTCCAATTGTCCTTG | Phage Genome Fragment Generation |
| 21099R | ggctaccgtctctACGATAGCCTCTTGGAG | Phage Genome Fragment Generation |
| 21096F | ggctaccgtctctTCGTCAAGATGTCCATGATTAG | Phage Genome Fragment Generation |
| 25539R | ggctaccgtctcgAAGGAGTCAATCTGTTGAC | Phage Genome Fragment Generation |
| 25536F | ggctaccgtctcgCCTTCACGACTAAAGATGG | Phage Genome Fragment Generation |
| 31046R | ggctaccgtctcgGCCATAGGTGACTTAGCAACATC | Phage Genome Fragment |

| | | |
|---|---|---|
| | | Generation |
| 31043F | ggctaccgtctcgTGGCTGGACAGTTGGAAAC | Phage Genome Fragment Generation |
| 35522R | ggctaccgtctcgGTGGTCATGTAAAGGCGCC | Phage Genome Fragment Generation |
| 35519F | ggctaccgtctcgCCACAAACGGTTTTGACTGTGG | Phage Genome Fragment Generation |
| 39754R | ggctaccgtctcgTATTTTAAAGGACCCTATAGGAACTCC | Phage Genome Fragment Generation |
| 12196R | ggctaccgtctcgGCTTCGTCCATATCGAACATTAAGATAATC | delDNAP Genome Fragment Generation |
| 12193F | ggctaccgtctcgAAGCAGGGCGCAAAGCA | delDNAP Genome Fragment Generation |
| 13338R | GgctaccgtctcgTAGTGAGTCGTATTAGAATGGGACTCTCC | delDNAP Genome Fragment Generation |
| 13335F | ggctaccgtctcgACTAAAGGAGACACACCATG | delDNAP Genome Fragment Generation |
| 16618R | ggctaccgtctcgATTTACCATCCTTACATACAACC | delDNAP Genome Fragment Generation |
| 16615F | ggctaccgtctcgAAATTTAGTAAGGTTCAGGTTAAAACAGCC | delDNAP Genome Fragment Generation |
| 17736R | ggctaccgtctcgGCACCTGCCCAAGCCTTC | delDNAP Genome Fragment Generation |
| T7 pol Amino + 9 His + vector | TGCATCACCATCACCATCACCATCACCATATGATCGTTTCTGACATC | PolA/TrxA cloning |
| T7 pol Carboxy + interlude | ATGTATATCTCCTGTTTAAACTCAGTGGCAAATCGCCCAATT | PolA/TrxA cloning |
| TrxA amino + interlude | GTTTAAACAGGAGATATACATATGAGCGATAAAATTATTCAC | PolA/TrxA cloning |
| TrxA carboxy + vector | GTGGCGGCCGCTCTATTATTACGCCAGGTTAGCGTCGAGGAA | PolA/TrxA cloning |
| Right vector + TrxA carboxy | CTCGACGCTAACCTGGCGTAATAATAGAGCGGCCGCCACCG | PolA/TrxA cloning |
| Left vector + T7 amino + 9 His | AGAAACGATCATATGGTGATGGTGATGGTGATGGTGATGCATATGTATATCTCC | PolA/TrxA cloning |
| 15660F | CCTTGCGCAAATTCCGGG | Sanger sequencing |
| 16202R | GCAGATTGCAGTAGGGTATTCA | Sanger sequencing |
| RNAP_F | GTTCAGGACATCTACGGGATTGTTGCTAA | qPCR RNA polymerase |
| RNAP_R | TCTCATCGGTCACGGTAACTACTTCGT | qPCR RNA polymerase |
| Term_F | AACCTTAGTGATGCCGAGAAGTACCC | qPCR Terminase large subunit |
| Term_R | TTCGGAAGCCACTGGTAATGCATTG | qPCR Terminase large subunit |

*Uppercase bases denote the portion of the primer with homology to T7 gDNA and used in annealing temperature calculation. Lowercase bases are BsmBI adapters

**Mutational primers

Supplementary Table S2. Genome fragment break points and PCR conditions

| Fragment | Start (nucleotide position)† | Stop (nucleotide position)† | FWD Primer Name | REV Primer Name | Annealing Temp (°C) | Fragment Length (bp) | Length with Extension (bp) |
|---|---|---|---|---|---|---|---|
| F12-1c | 39751 | 874 | 39751F | 874R | 61 | 901 | 927 |
| F12-2 | 871 | 1998 | 871F | 1998R | 65 | 1128 | 1154 |
| F12-3 | 1995 | 5534 | 1995F | 5534R | 64 | 3540 | 3566 |
| F12-4 | 5531 | 8857 | 5531F | 8857R | 66 | 3327 | 3353 |
| F12-5 | 8854 | 11208 | 8854F | 11208R | 63 | 2355 | 2381 |
| F12-6 | 11205 | 14405 | 11205F | 14405R | 64 | 3201 | 3227 |
| F12-7a | 14402 | 15940 | 14402F | 15940R* | 67 | 1539 | 1565 |
| F12-7b | 15937 | 17736 | 15937F | 17736R | 68 | 1800 | 1826 |
| F12-8 | 17733 | 21099 | 17733F | 21099R | 61 | 3367 | 3393 |
| F12-9 | 21096 | 25539 | 21096F | 25539R | 60 | 4444 | 4470 |
| F12-10 | 25536 | 31046 | 25536F | 31046R | 61 | 5511 | 5537 |
| F12-11 | 31043 | 35522 | 31043F | 35522R | 66 | 4480 | 4506 |
| F12-12 | 35519 | 39754 | 35519F | 39754R | 64 | 4236 | 4264 |
| **F12-7c | 12205 | 12196 | 11205F | 12196R | 66 | 992 | 1018 |
| **F12-7d | 12193 | 13338 | 12193F | 13338R | 69 | 1146 | 1172 |
| **F12-7e_sfGPF | 13335 | 16618 | 13335F | 16618R | 60 | 1886 | 1912 |
| **F12-7f | 16615 | 17736 | 16615F | 17736R | 66 | 1122 | 1148 |

† Positioning based on WT T7 genome (GenBank V01146)

* One of three 15940R primers (see Supplementary Table S1)

** Fragments used only in ΔDNAP genome assembly

Supplementary Table S3. Synthetic F12-7e_sfGPF Twist Fragment

| Fragment | Sequence |
|---|---|
| F12-7e_sfGPF | CCCAGTCACGACGTTGTAAAACGCGTCTCCACTAAAGGAGACACACCATGTTCAAACT<br>GATTAAGAAGTTAGGCCAACTGCTGGTTCGTATGTACAACGTGGAAGCCAAGCGACTG<br>AACGATGAGGCTCGTAAAGAGGCCACACAGTCACGCGCTCTGGCGATTCGCTCCAACG<br>AACTGGCTGACAGTGCATCCACTAAAGTTACCGAGGCTGCCCGTGTGGCAAACCAAGC<br>TCAACAGCTTTCCAAATTCTTTGAGTAATCAAACAGGAGAAACCATTATGTCTAACGT<br>AGCTGAAACTATCCGTCTATCCGATACAGCTGACCAGTGGAACCGTCGAGTCCACATC<br>AACGTTCGCAACGGTAAGGCGACTATGGTTTACCGCTGGAAGGACTCTAAGTCCTCTA<br>AGAATCACACTCAGCGTATGACGTTGACAGATGAGCAAGCACTGCGTCTGGTCAATGC<br>GCTTACCAAAGCTGCCGTGACAGCAATTCATGAAGCTGGTCGCGTCAATGAAGCTATG<br>GCTATCCTCGACAAGATTGATAACTAAGAGTGGTATCCTCAAGGTCGCCAAAGTGGTG<br>GCCTTCATGAATACTATTCGACTCACTATAGGAGATATTACCATGCGTGACCCTAAAG<br>TTATCCAAGCAGAAATCGCTAAACTGGAAGCTGAACTGGAGGACGTTAAGTACCATGA<br>AGCTAAGACTCGCTCCGCTGTTCACATCTTGAAGAACTTAGGCTGGACTTGGACAAGA<br>CAGACTGGCTGGAAGAAACCAGAAGTTACCAAGCTGAGTCATAAGGTGTTCGATAAGG<br>ACACTATGACCCACATCAAGGCTGGTGATTGGGTTAAGGTTGACATGGGAGTTGTTGG<br>TGGATACGGCTACGTCCGCTCAGTTAGTGGCAAATATGCACAAGTGTCATACATCACA<br>GGTGTTACTCCACGCGGTGCAATCGTTGCCGATAAGACCAACATGATTCACACAGGTT<br>TCTTGACAGTTGTTTCATATGAAGAGATTGTTAAGTCACGATAATCAATAGGAGAAAT<br>CAATATGAGCAAAGGAGAAGAACTTTTCACTGGAGTTGTCCCAATTCTTGTTGAATTA<br>GATGGTGATGTTAATGGGCACAAATTTTCTGTCCGTGGAGAGGGTGAAGGTGATGCTA<br>CAAACGGAAAACTCACCCTTAAATTTATTTGCACTACTGGAAAACTACCTGTTCCATG<br>GCCAACACTTGTCACTACTCTGACCTATGGTGTTCAATGCTTTTCCCGTTATCCGGAT<br>CACATGAAACGGCATGACTTTTTCAAGAGTGCCATGCCCGAAGGTTATGTACAGGAAC<br>GCACTATATCTTTCAAAGATGACGGGACCTACAAGACGCGTGCTGAAGTCAAGTTTGA<br>AGGTGATACCCTTGTTAATCGTATCGAGTTAAAAGGTATTGATTTTAAAGAAGATGGA<br>AACATTCTCGGACACAAACTGGAGTACAACTTTAACTCACACAATGTATACATCACGG<br>CAGACAAACAAAAGAATGGAATCAAAGCTAACTTCAAAATTCGCCACAACGTTGAAGA<br>TGGTTCCGTTCAACTAGCAGACCATTATCAACAAAATACTCCAATTGGTGATGGCCCT<br>GTCCTTTTACCAGACAACCATTACCTGTCGACACAATCTGTCCTTTCGAAAGATCCCA<br>ACGAAAAGCGTGACCACATGGTCCTTCTTGAGTTTGTAACTGCTGCTGGGATTACACA<br>TGGCATGGATGAGCTCTACAAATGATACAGGAGGCTACTCATGAACGAAAGACACTTA<br>ACAGGTGCTGCTTCTGAAATGCTAGTAGCCTACAAATTTACCAAAGCTGGGTACACTG<br>TCTATTACCCTATGCTGACTCAGAGTAAAGAGGACTTGGTTGTATGTAAGGATGGTAA<br>ATAGAGACGCCTGTGTGAAATTGTTATCCGCT |

### Supplementary Files

#### Supplementary File S1. Codon-shuffled plasmid sequence

```
LOCUS       T7_DNA_Pol_pSL003        7308 bp    DNA     circular UNA 14-MAY-2026
DEFINITION  Generated by Golden Gate/Type IIS Cloning (delLacO_pSL003_pET28a,
            T7 DNAP codon shuffled A, T7 DNAP codon shuffled B).
ACCESSION   urn.local...11n9-km7ppin
VERSION     urn.local...11n9-km7ppin
KEYWORDS    .
SOURCE
  ORGANISM  .
FEATURES             Location/Qualifiers
     primer_bind     303..322
                     /Sequence="TAATACGACTCACTATAGGG"
                     /Tm="50.3"
                     /%GC="40.0"
                     /Hairpin_Tm="None"
                     /Self_Dimer_Tm="4.4"
                     /label="T7"
     promoter        303..321
                     /note="promoter for bacteriophage T7 RNA polymerase"
                     /standard_name="T7 promoter"
     RBS             344..366
     CDS             373..2487
                     /label="T7 PolA Codon Shuffled CDS"
     primer_bind     complement(2545..2563)
                     /Sequence="GCTAGTTATTGCTCAGCGG"
                     /Tm="56.2"
                     /%GC="52.6"
                     /Hairpin_Tm="None"
                     /Self_Dimer_Tm="None"
                     /label="T7 Term"
     terminator      2559..2606
                     /note="transcription terminator for bacteriophage T7 RNA
                     polymerase"
                     /standard_name="T7 terminator"
     rep_origin      2643..3098
                     /label="f1 ori"
     CDS             complement(3190..4005)
                     /codon_start=1
                     /gene="<i>aph(3')-Ia</i>"
                     /note="confers resistance to kanamycin"
                     /product="aminoglycoside phosphotransferase"
                     /transl_table=1
                     /translation="MSHIQRETSCSRPRLNSNMDADLYGYKWARDNVGQSGATIYRLY
                     GKPDAPELFLKHGKGSVANDVTDEMVRLNWLTEFMPLPTIKHFIRTPDDAWLLTTAIP
                     GKTAFQVLEEYPDSGENIVDALAVFLRRLHSIPVCNCPFNSDRVFRLAQAQSRMNNGL
                     VDASDFDDERNGWPVEQVWKEMHKLLPFSPDSVVTHGDFSLDNLIFDEGKLIGCIDVG
                     RVGIADRYQDLAILWNCLGEFSPSLQKRLFQKYGIDNPDMNKLQFHLMLDEFF*"
                     /standard_name="KanR"
     rep_origin      4127..4715
                     /label="ori"
     CDS             complement(5142..5333)
                     /codon_start=1
```

/gene="<i>rop</i>"
/product="Rop protein"
/transl_table=1
/translation="VTKQEKTALNMARFIRSQTLTLLEKLNELDADEQADICESLHDH
ADELYRSCLARFGDDGENL*"
/standard_name="rop"
protein_bind complement(6108..6129)
/label="CAP binding site"
CDS complement(6142..7224)
/codon_start=1
/gene="<i>lacI</i>"
/note="The <i>lac</i> repressor binds to the <i>lac</i>
operator to inhibit transcription in <i>E. coli. </i>This
inhibition can be relieved by adding lactose or
isopropyl-β-D-thiogalactopyranoside (IPTG)."
/product="<i>lac</i> repressor"
/transl_table=1
/translation="VKPVTLYDVAEYAGVSYQTVSRVVNQASHVSAKTREKVEAAMAE
LNYIPNRVAQQLAGKQSLLIGVATSSLALHAPSQIVAAIKSRADQLGASVVVSMVERS
GVEACKAAVHNLLAQRVSGLIINYPLDDQDAIAVEAACTNVPALFLDVSDQTPINSII
FSHEDGTRLGVEHLVALGHQQIALLAGPLSSVSARLRLAGWHKYLTRNQIQPIAEREG
DWSAMSGFQQTMQMLNEGIVPTAMLVANDQMALGAMRAITESGLRVGADISVVGYDDT
EDSSCYIPPLTTIKQDFRLLGQTSVDRLLQLSQGQAVKGNQLLPVSLVKRKTTLAPNT
QTASPRALADSLMQLARQVSRLESGQ*"
/standard_name="lacI"
promoter complement(7225..7302)
/gene="<i>lacI</i>"
/note="<br>"
/standard_name="lacI promoter"
ORIGIN
1 ctcgacgctc tcccttatgc gactcctgca ttaggaagca gcccagtagt aggttgaggc
61 cgttgagcac cgccgccgca aggaatggtg catgcaagga gatggcgccc aacagtcccc
121 cggccacggg gcctgccacc atacccacgc cgaaacaagc gctcatgagc ccgaagtggc
181 gagcccgatc ttccccatcg gtgatgtcgg cgatataggc gccagcaacc gcacctgtgg
241 cgccggtgat gccggccacg atgcgtccgg cgtagaggat cgagatctcg atcccgcgaa
301 attaatacga ctcactatag gggaattccc ctctagaaat aattttgttt aactttaaga
361 aggagatata ccatgattgt gtccgatatt gaggcgaatg ctttgctgga atccgtgacc
421 aagtttcatt gtggagtcat ttatgattat agcactgcag aatatgtctc ctatcgcccc
481 tccgatttcg gcgcatactt ggacgcactt gaagctgaag tcgcgcgcgg aggcctgatc
541 gtattccata acggccataa atacgatgtg ccagcgctga cgaagcttgc caaacttcag
601 ttaaatcgtg aattccattt accccgcgaa aattgcatcg atacgctagt tttaagccgt
661 cttatccaca gtaatttaaa agatacggat atgggcttac tgcgcagtgg gaaacttccg
721 ggcaagcgtt tcggtagcca tgcgcttgaa gcatggggct accgtcttgg tgagatgaaa
781 ggcgagtata aggatgattt caaacgcatg ttagaggaac aaggagagga gtatgtggat
841 ggtatggaat ggtggaattt taatgaggaa atgatggatt acaatgtcca agatgtcgta
901 gtgacaaagg cgttgttaga aaaattgctg agtgataagc actattttcc cccagaaatc
961 gatttcaccg atgttggtta tacaaccttt tggagtgaga gtttagaagc ggtggatatc
1021 gagcaccgcg cagcgtggtt attggcgaag caggaacgta atggctttcc cttcgatacg
1081 aaggccattg aggaactata tgtggaactt gcagcgcgtc gtagcgaact tttacgcaag
1141 ctcacggaga catttgggtc ttggtaccaa cccaagggag ggacggaaat gttttgtcac
1201 ccacgtacgg ggaaaccttt gccaaagtat ccccgcatca aaaccccaaa ggtaggcgga
1261 attttcaaaa agcccaaaaa taaggcgcaa cgtgagggac gcgaaccgtg cgagttggac
1321 acgcgtgaat atgtggcagg ggccccctat acgcccgtag agcacgtagt tttcaatcca
1381 agcagtcgcg atcatatcca aaaaaagtta caggaagccg gatgggtgcc cactaaatat
1441 acggacaaag gcgcgcccgt cgtcgatgac gaagtcctgg agggcgtccg cgttgacgat
1501 ccagaaaaac aggctgccat tgatctgatc aaggaatacc tgatgatcca aaaacgcatt
1561 ggccaaagcg cggaaggtga taaggcctgg ttacgctacg tggcagaaga cggcaaaatc

1621 cacggcagtg tcaatcccaa cggggccgtg accggacgcg caacacacgc ctttccgaat
1681 ttggcacaga tccccggagt gcgtagtccc tacggggaac aatgccgtgc agccttcggt
1741 gcagaacatc accttgacgg tattaccggc aaaccatggg tccaagcagg tattgatgcg
1801 agcgggttag aattgcgctg tcttgcacat tttatggccc gtttcgacaa tggtgaatat
1861 gcacatgaaa tcttgaatgg ggatattcat accaaaaatc aaattgccgc ggagttaccg
1921 acgcgcgaca atgcaaaaac ttttatttac ggctttttgt acggcgcggg ggacgaaaag
1981 atcggtcaaa tcgtgggtgc cggaaaggaa cgtggcaaag agttgaagaa gaagtttctg
2041 gaaaatacac cggcaatcgc ggccttacgt gaatcaattc agcaaacttt ggttgaaagt
2101 agccagtggg tggcaggggа acagcaggtt aaatggaagc gtcgttggat taagggctta
2161 gacggccgca aagtccatgt gcgctcaccg catgcagcgc tgaacacgtt gctacagagt
2221 gccggggcat taatttgtaa gttatggatc attaaaacgg aggaaatgtt agtggaaaag
2281 ggtcttaaac acggatggga cggcgatttc gcttatatgg cctgggttca cgacgagatt
2341 caggtcggat gtcgcacaga ggaaatcgcc caagtcgtga tcgaaactgc ccaggaggct
2401 atgcgttggg taggcgatca ttggaatttt cgctgcttgc ttgacacgga gggaaaaatg
2461 gggccgaact gggcaatctg tcattgattc tgtaagatcc ggctgctaac aaagcccgaa
2521 aggaagctga gttggctgct gccaccgctg agcaataact agcataaccc cttggggcct
2581 ctaaacgggt cttgaggggt tttttgctga aaggaggaac tatatccgga ttggcgaatg
2641 ggacgcgccc tgtagcggcg cattaagcgc ggcgggtgtg gtggttacgc gcagcgtgac
2701 cgctacactt gccagcgccc tagcgcccgc tcctttcgct ttcttccctt cctttctcgc
2761 cacgttcgcc ggctttcccc gtcaagctct aaatcggggg ctccctttag ggttccgatt
2821 tagtgcttta cggcacctcg accccaaaaa acttgattag ggtgatggtt cacgtagtgg
2881 gccatcgccc tgatagacgg tttttcgccc tttgacgttg gagtccacgt tctttaatag
2941 tggactcttg ttccaaactg gaacaacact caaccctatc tcggtctatt cttttgattt
3001 ataagggatt ttgccgattt cggcctattg gttaaaaaat gagctgattt aacaaaaatt
3061 taacgcgaat tttaacaaaa tattaacgct tacaatttag gtggcacttt tcggggaaat
3121 gtgcgcggaa cccctatttg tttatttttc taaatacatt caaatatgta tccgctcatg
3181 aattaattct tagaaaaact catcgagcat caaatgaaac tgcaatttat tcatatcagg
3241 attatcaata ccatattttt gaaaaagccg tttctgtaat gaaggagaaa actcaccgag
3301 gcagttccat aggatggcaa gatcctggta tcggtctgcg attccgactc gtccaacatc
3361 aatacaacct attaatttcc cctcgtcaaa aataaggtta tcaagtgaga aatcaccatg
3421 agtgacgact gaatccggtg agaatggcaa aagtttatgc atttctttcc agacttgttc
3481 aacaggccag ccattacgct cgtcatcaaa atcactcgca tcaaccaaac cgttattcat
3541 tcgtgattgc gcctgagcga gacgaaatac gcgatcgctg ttaaaaggac aattacaaac
3601 aggaatcgaa tgcaaccggc gcaggaacac tgccagcgca tcaacaatat tttcacctga
3661 atcaggatat tcttctaata cctggaatgc tgttttcccg gggatcgcag tggtgagtaa
3721 ccatgcatca tcaggagtac ggataaaatg cttgatggtc ggaagaggca taaattccgt
3781 cagccagttt agtctgacca tctcatctgt aacatcattg gcaacgctac ctttgccatg
3841 tttcagaaac aactctggcg catcgggctt cccatacaat cgatagattg tcgcacctga
3901 ttgcccgaca ttatcgcgag cccatttata cccatataaa tcagcatcca tgttggaatt
3961 taatcgcggc ctagagcaag acgtttcccg ttgaatatgg ctcataacac cccttgtatt
4021 actgtttatg taagcagaca gttttattgt tcatgaccaa aatcccttaa cgtgagtttt
4081 cgttccactg agcgtcagac cccgtagaaa agatcaaagg atcttcttga gatccttttt
4141 ttctgcgcgt aatctgctgc ttgcaaacaa aaaaaccacc gctaccagcg gtggtttgtt
4201 tgccggatca agagctacca actctttttc cgaaggtaac tggcttcagc agagcgcaga
4261 taccaaatac tgtccttcta gtgtagccgt agttaggcca ccacttcaag aactctgtag
4321 caccgcctac atacctcgct ctgctaatcc tgttaccagt ggctgctgcc agtggcgata
4381 agtcgtgtct taccgggttg gactcaagac gatagttacc ggataaggcg cagcggtcgg
4441 gctgaacggg gggttcgtgc acacagccca gcttggagcg aacgacctac accgaactga
4501 gatacctaca gcgtgagcta tgagaaagcg ccacgcttcc cgaagggaga aaggcggaca
4561 ggtatccggt aagcggcagg gtcggaacag gagagcgcac gagggagctt ccagggggaa
4621 acgcctggta tctttatagt cctgtcgggt ttcgccacct ctgacttgag cgtcgatttt
4681 tgtgatgctc gtcagggggg cggagcctat ggaaaaacgc cagcaacgcg gcctttttac
4741 ggttcctggc cttttgctgg ccttttgctc acatgttctt tcctgcgtta tcccctgatt
4801 ctgtggataa ccgtattacc gcctttgagt gagctgatac cgctcgccgc agccgaacga
4861 ccgagcgcag cgagtcagtg agcgaggaag cggaagagcg cctgatgcgg tattttctcc
4921 ttacgcatct gtgcggtatt tcacaccgca atggtgcact ctcagtacaa tctgctctga
4981 tgccgcatag ttaagccagt atacactccg ctatcgctac gtgactgggt catggctgcg

5041 ccccgacacc cgccaacacc cgctgacgcg ccctgacggg cttgtctgct cccggcatcc
5101 gcttacagac aagctgtgac cgtctccggg agctgcatgt gtcagaggtt ttcaccgtca
5161 tcaccgaaac gcgcgaggca gctgcggtaa agctcatcag cgtggtcgtg aagcgattca
5221 cagatgtctg cctgttcatc cgcgtccagc tcgttgagtt tctccagaag cgttaatgtc
5281 tggcttctga taaagcgggc catgttaagg gcggtttttt cctgtttggt cactgatgcc
5341 tccgtgtaag ggggatttct gttcatgggg gtaatgatac cgatgaaacg agagaggatg
5401 ctcacgatac gggttactga tgatgaacat gcccggttac tggaacgttg tgagggtaaa
5461 caactggcgg tatggatgcg gcgggaccag agaaaaatca ctcagggtca atgccagcgc
5521 ttcgttaata cagatgtagg tgttccacag ggtagccagc agcatcctgc gatgcagatc
5581 cggaacataa tggtgcaggg cgctgacttc cgcgtttcca gactttacga aacacggaaa
5641 ccgaagacca ttcatgttgt tgctcaggtc gcagacgttt tgcagcagca gtcgcttcac
5701 gttcgctcgc gtatcggtga ttcattctgc taaccagtaa ggcaaccccg ccagcctagc
5761 cgggtcctca acgacaggag cacgatcatg cgcacccgtg gggccgccat gccggcgata
5821 atggcctgct tctcgccgaa acgtttggtg gcgggaccag tgacgaaggc ttgagcgagg
5881 gcgtgcaaga ttccgaatac cgcaagcgac aggccgatca tcgtcgcgct ccagcgaaag
5941 cggtcctcgc cgaaaatgac ccagagcgct gccggcacct gtcctacgag ttgcatgata
6001 aagaagacag tcataagtgc ggcgacgata gtcatgcccc gcgcccaccg gaaggagctg
6061 actgggttga aggctctcaa gggcatcggt cgagatcccg gtgcctaatg agtgagctaa
6121 cttacattaa ttgcgttgcg ctcactgccc gctttccagt cgggaaacct gtcgtgccag
6181 ctgcattaat gaatcggcca acgcgcgggg agaggcggtt tgcgtattgg gcgccagggt
6241 ggtttttctt ttcaccagtg agacgggcaa cagctgattg cccttcaccg cctggccctg
6301 agagagttgc agcaagcggt ccacgctggt ttgccccagc aggcgaaaat cctgtttgat
6361 ggtggttaac ggcgggatat aacatgagct gtcttcggta tcgtcgtatc ccactaccga
6421 gatatccgca ccaacgcgca gcccggactc ggtaatggcg cgcattgcgc ccagcgccat
6481 ctgatcgttg gcaaccagca tcgcagtggg aacgatgccc tcattcagca tttgcatggt
6541 ttgttgaaaa ccggacatgg cactccagtc gccttcccgt tccgctatcg gctgaatttg
6601 attgcgagtg agatatttat gccagccagc cagacgcaga cgcgccgaga cagaacttaa
6661 tgggcccgct aacagcgcga tttgctggtg acccaatgcg accagatgct ccacgcccag
6721 tcgcgtaccg tcttcatggg agaaaataat actgttgatg ggtgtctggt cagagacatc
6781 aagaaataac gccggaacat tagtgcaggc agcttccaca gcaatggcat cctggtcatc
6841 cagcggatag ttaatgatca gcccactgac gcgttgcgcg agaagattgt gcaccgccgc
6901 tttacaggct tcgacgccgc ttcgttctac catcgacacc accacgctgg cacccagttg
6961 atcggcgcga gatttaatcg ccgcgacaat ttgcgacggc gcgtgcaggg ccagactgga
7021 ggtggcaacg ccaatcagca acgactgttt gcccgccagt tgttgtgcca cgcggttggg
7081 aatgtaattc agctccgcca tcgccgcttc cactttttcc cgcgttttcg cagaaacgtg
7141 gctggcctgg ttcaccacgc gggaaacggt ctgataagag acaccggcat actctgcgac
7201 atcgtataac gttactggtt tcacattcac caccctgaat tgactctctt ccgggcgcta
7261 tcatgccata ccgcgaaagg ttttgcgcca ttcgatggtg tccgggat

//
